# GAPDH controls extracellular vesicle biogenesis and enhances therapeutic potential of EVs in silencing the Huntingtin gene in mice via siRNA delivery

**DOI:** 10.1101/2020.01.09.899880

**Authors:** Ghulam Hassan Dar, Cláudia C. Mendes, Wei-Li Kuan, Mariana Conceição, Samir El-Andaloussi, Imre Mager, Thomas C. Roberts, Roger A. Barker, Deborah C. I. Goberdhan, Clive Wilson, Matthew J.A. Wood

**Affiliations:** Department of Paediatrics, University of Oxford, OX1 3QX, Oxford, UK; Department of Physiology, Anatomy and Genetics, University of Oxford, OX1 3QX, Oxford, UK; Centre for Brain Repair, Department of Clinical Neurosciences, University of Cambridge, CB2 0PY, Cambridge, UK; Department of Laboratory Medicine, Karolinska Institute, 14186, Sweden; MDUK Oxford Neuromuscular Centre, University of Oxford, OX2 9DU, Oxford, UK

## Abstract

Extracellular vesicles (EVs) are biological nanoparticles with important roles in intercellular communication and pathophysiology. Their capacity to transfer biomolecules between cells has sparked efforts to bioengineer EVs as drug delivery vehicles. However, a better understanding of EV biogenesis mechanisms and function is required to unleash their considerable therapeutic potential. Here we demonstrate a novel role for GAPDH, a glycolytic enzyme, in EV assembly and secretion, and we exploit these findings to develop a GAPDH-based methodology to load therapeutic siRNAs onto EVs for targeted drug delivery to the brain. In a series of experiments, we observe high levels of GAPDH binding to the outer surface of EVs *via* a phosphatidylserine binding motif, designated as G58, and discover that the tetrameric nature of GAPDH promotes extensive EV aggregation. Studies in a *Drosophila* EV biogenesis model demonstrate that GAPDH is absolutely required for normal generation of intraluminal vesicles in endosomal compartments and promotes vesicle clustering both inside and outside the cell. Fusing a GAPDH-derived G58 peptide to dsRNA-binding motifs permits highly efficient loading of RNA-based drugs such as siRNA onto the surface of EVs. Such vesicles efficiently deliver siRNA to target cells *in vitro* and into the brain of a Huntington’s disease mouse model after systemic injection, resulting in silencing of the huntingtin gene in multiple anatomical regions of the brain and modulation of phenotypic features of disease. Taken together, our study demonstrates a novel role for GAPDH in EV biogenesis, and that the presence of free GAPDH binding sites on EVs can be effectively exploited to substantially enhance the therapeutic potential of EV-mediated drug delivery to the brain.

---

During the last decade, a new paradigm of cell-to-cell communication has emerged involving extracellular vesicles (EVs), with implications for both normal and pathological physiology^1,2^. There are several classes of EVs, but small EVs with a diameter of 30-150 nm are the best studied^3^. They are secreted by cells as exosomes formed in intracellular endosomal compartments or as microvesicles shed from the cell surface. Given their unique biological and pharmacological characteristics, EVs have attracted tremendous interest as mediators of physiological and pathophysiological processes, and as vehicles for drug delivery^4^. Immunological inertness, and ability to cross biological barriers and carry bioactive molecules are attractive features of EVs that can be exploited in drug delivery and therapeutic applications^5,6^. However, lack of efficient drug loading methods, and an incomplete understanding of EV biogenesis and uptake mechanisms remain critical challenges that need to be addressed^7,8^. Current methods of loading therapeutic molecules into EVs such as electroporation, genetic engineering of host cells and chemical conjugation, are limited by low efficiency, toxicity and lack of scalability^9,10^. Moreover, they produce a heterogeneous population of EVs that imposes further complexity in understanding the phenotypic effects of EVs in target cells^11,12^. It is important to understand the intracellular pathways of EV biogenesis, so that the natural characteristics of EVs can be exploited for therapeutic applications.

In a set of experiments designed to express the N-terminal region of lactoferrin (designated as lactoferrin N) on the surface of EVs, we observed that lactoferrin N, which had been fused to DC-LAMP (Dendritic cell lysosomal associated membrane glycoprotein) to attach it to the surface of EVs, was specifically cleaved from this anchor. Surprisingly, cleaved lactoferrin N was still found associated with the EV surface (**Supporting data, Fig. 1 & 2**). Previously, it was demonstrated that cells take up extracellular iron by secreting EVs carrying surface glyceraldehyde-3-phosphate dehydrogenase (GAPDH), which attaches to the iron-binding proteins lactoferrin and transferrin^13^. We confirmed the presence of GAPDH on the outside of EVs isolated from different cell sources using a protease digestion assay **(Extended Fig. 1a & 1b)**. An enzyme activity assay indicated that this surface GAPDH was enzymatically active **(Extended Fig. 1c**). Co-immunoprecipitation experiments of HEK293T cell lysates and EVs demonstrated that the lactoferrin N domain is physically associated with GAPDH in cells and on the surface of EVs **(Extended Fig. 1d)**. Furthermore, incubation of isolated EVs with purified lactoferrin N protein resulted in efficient binding of the protein to the EV surface. However, incubating purified EVs with a different domain of lactoferrin, the N1.1 domain, which lacks the GAPDH binding motif, did not result in binding to the EV surface (**Extended Fig. 1e & Extended Fig. 2a**). Taken together, these experiments demonstrate the role of GAPDH in tethering lactoferrin N on the surface of EVs.

To enhance loading of lactoferrin N on to EVs, we increased the GAPDH concentration on the EV surface. Incubation of isolated EVs with GAPDH protein resulted in extensive surface binding of GAPDH, as confirmed by immunoblotting, UV-spectrophotometry and GAPDH enzymatic activity **(Fig 1a, 1b & Extended Fig. 2c)**. Binding of GAPDH to EVs occurred for all cell sources tested with or without the presence of serum proteins **(Extended Fig. 2b-2d)**. Interestingly, incubation of EVs with GAPDH resulted in an increase in EV particle size, as determined by nanoparticle tracking analysis **(Fig 1c)**. Electron microscopy of HEK293T EVs after incubation with GAPDH revealed the formation of long branched chains of EVs **(Fig. 1d)**. EVs derived from HeLa, B16 F10 and mesenchymal stem cells formed conspicuous thread-like structures when incubated with GAPDH protein, suggesting that GAPDH induces EV aggregation (**Fig. 1e)**.

**Figure 1.**
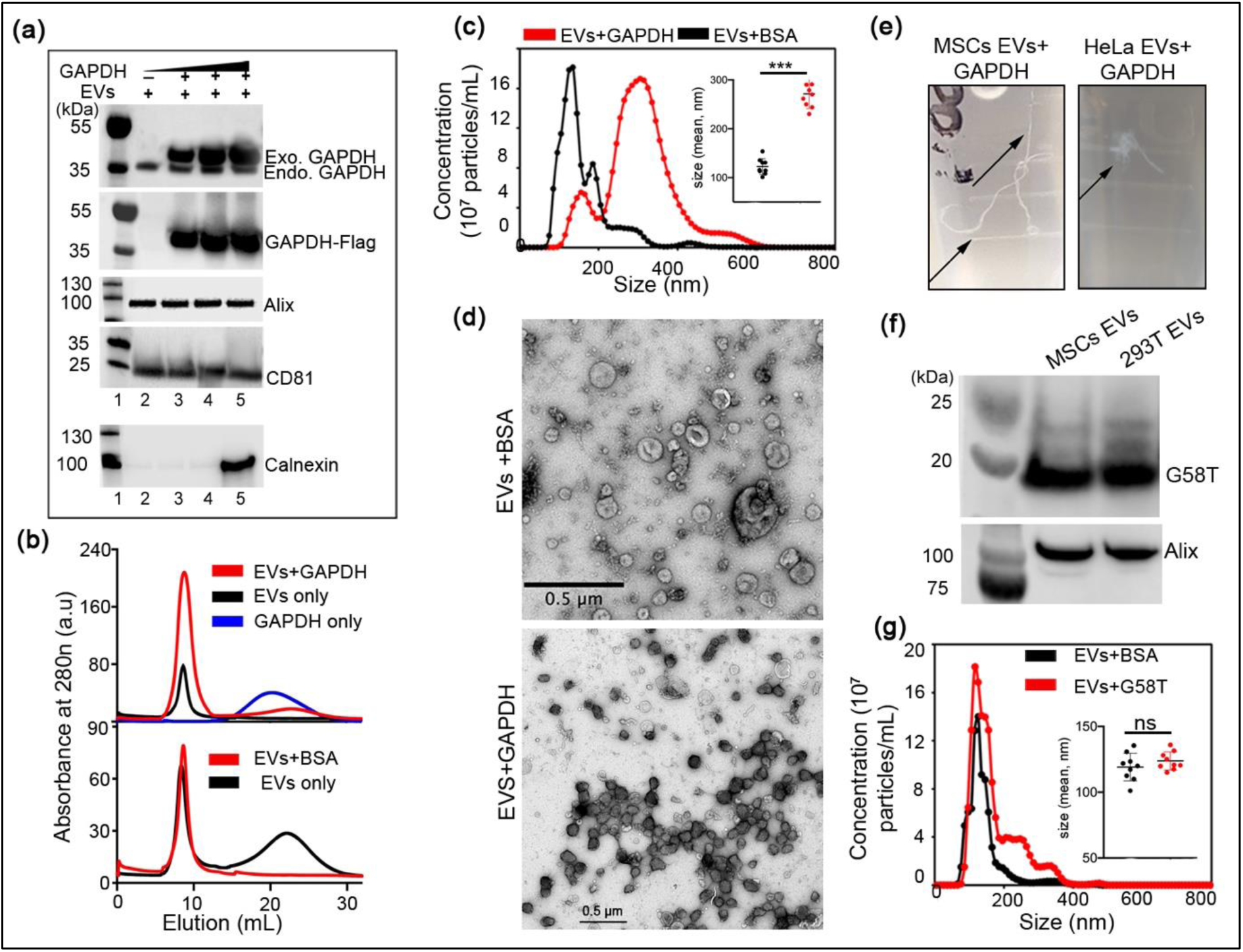
Surface binding of GAPDH leads to aggregation of EVs: **(a)** Western blot showing exogenous binding of GAPDH to HEK293T EVs. Increasing concentrations of histidine-(His6) and Flag-tagged GAPDH (lane 3,4 and 5) were incubated with a fixed number of EVs (see details in methods). Endogenous GAPDH (Endo. GAPDH) present naturally on the EV surface is shown in the top blot along with the exogenous GAPDH (Exo. GAPDH). Alix and CD81 are EV protein markers used as a positive control. Calnexin (bottom blot) was used to demonstrate the purity of EV samples. In this blot, lane 2, 3 and 4 represent EVs, and lane 5 represents cell lysate. Representative blots (n > 3). **(b)** UV-absorbance spectrum of EVs after passing through gel-filtration column. Increase in the absorbance of EVs+GAPDH peak indicates binding of GAPDH protein. Purified EVs were used for incubation with either GAPDH (upper chromatogram) or BSA protein (lower chromatogram). The first peak (at ∼10 ml elution) represents EVs and the second peak (around 20 ml elution) represents unbound proteins. Representative graphs (n > 3). **(c)** NTA profile showing the size distribution of purified HEK293T EVs after incubation with either GAPDH or BSA proteins respectively. Binding of GAPDH to the EVs, shifts their size. Inset is the scatter plot representing size (mean) of EVs (red; EVs+GAPDH, black; EVs+BSA). Data shown as mean ± s.d, n=9, ***p< 0.0001 when compared to EVs+BSA (two tailed t-test) **(d)** Electron microscopy images of EVs incubated with either BSA or GAPDH protein. Excess of unbound GAPDH protein was removed by gel-filtration and Uranyl Oxalate, pH 7 was used to stain the EVs. Images were collected at 120kVon FEI Tecnai 12 TEM with Gatan one view CMOS Camera. Representative images (n = 2). (Raw images are in Supporting data, Fig. 3a). **(e)** Photographic images (taken with Canon EOS 200D) of tubes showing the formation of thread-like structures from MSCs and HeLa EVs after incubation with GAPDH for 2h at 4°C. Representative images (n = 2). (Raw images are in Supporting data, Fig. 3b). **(f)** Western blot showing binding of G58 peptide to HEK293T (designated as 293T) and MSC EVs. Second domain of TRBP protein was attached to G58 peptide for detection by TRBP2 antibody. Representative blots (n>4). **(g)** NTA profile showing the size distribution of HEK293T EVs after binding to the G58T protein. Inset is the scatter plot representing size (mean) of EVs after incubating with either BSA (black) or G58T protein (red). Data shown as mean ± s.d, n=9 (ns = non-significant, two tailed t-test). Binding of G58T protein to EVs did not significantly alter the size of EVs.

GAPDH isolated from rabbit brain tissues has been reported to induce fusion of synthetic lipid vesicles that contained phosphatidylserine (PS), cholesterol and plasmenylethanolamine^14^. The fusogenic activity of GAPDH can also play an important role in nuclear membrane formation^15,16^. GAPDH protein was found to bind to the nuclear membrane via a PS-binding domain, designated as G58, located between amino acids 70-90^17^. To determine whether the PS-binding domain of GAPDH is also responsible for mediating binding of GAPDH to the outer surface of EVs, we incubated purified G58 peptide with EVs for 2h at 4 °C and then separated EVs from the unbound G58 peptide using gel-filtration chromatography. Binding of G58 peptide to EVs was assessed by western blotting, which revealed extensive binding of the peptide to the EV surface **(Fig. 1f)**. Quantification of G58 binding on MSC- and HEK293T-derived EVs revealed approximately 1200 and 1400 G58 peptide binding sites per EV respectively **(Extended Fig. 3a-c)**. Moreover, binding of G58 did not significantly alter the size of EVs, suggesting that the tetrameric nature of GAPDH is responsible for aggregation of EVs **(Fig. 1g)**.

To assess the physiological role of GAPDH in EV biology, we exploited a *Drosophila melanogaster* model of exosome biogenesis. In secondary cells (SCs) of the *Drosophila* accessory gland (AG), exosomes are formed as intraluminal vesicles (ILVs) in highly enlarged Rab11-positive compartments and then secreted into the lumen of the AG, a storage site for seminal fluid^18,19^. These ILVs can be selectively marked by fluorescent transmembrane markers, such as a GFP-tagged form of the FGF receptor, Breathless (Btl-GFP). In SCs, ILVs form in clusters that surround a large dense-core granule (DCG) of aggregated protein and extend out to the limiting membrane of the Rab11 compartments^19^. *Drosophila* GAPDH is highly conserved and has similar EV binding properties to human GAPDH (**Extended Fig. 3d & 3e)**. Overexpression of human GAPDH specifically in adult SCs produced larger clusters of Btl-GFP-positive ILVs, an effect also observed when ILVs were marked with a second GFP-tagged transmembrane exosome marker, human CD63 **(Fig. 2b, Extended Fig. 4b)**. There was also increased clustering of Btl-GFP and CD63-GFP puncta in the AG lumen, suggesting that human GAPDH promotes vesicle aggregation *in vivo* **(Fig. 2c, Extended Fig. 4b)**.

**Figure 2:**
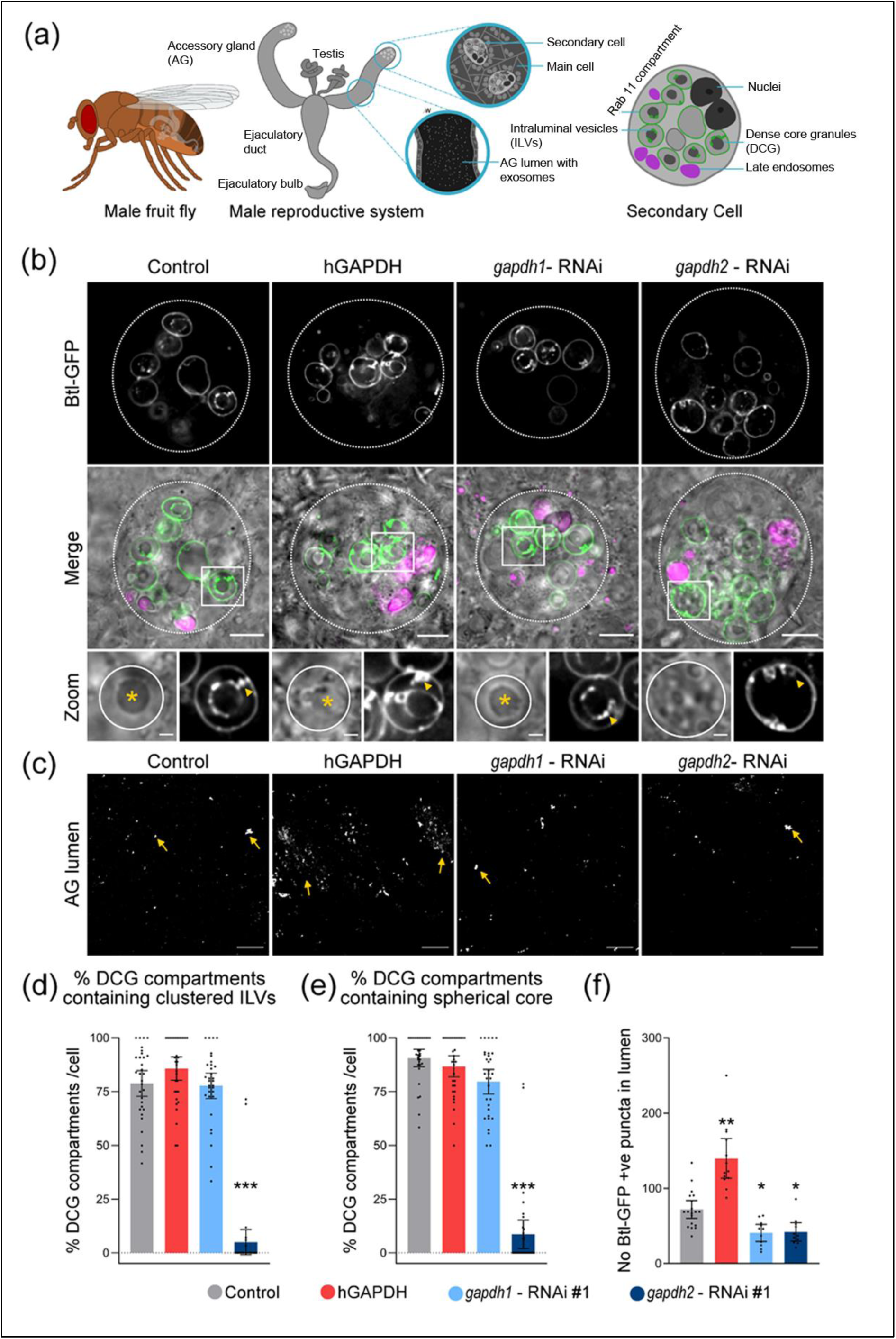
GAPDH regulates exosome biogenesis and clustering in *Drosophila* secondary cells: **(a)** Schematics show male fruit fly and its accessory gland (AG) containing main cells and secondary cells (SCs), which are only found at the distal tip of the gland. Exosomes can be visualised at the AG lumen as fluorescent puncta. A schematic of a secondary cell expressing a GFP-tagged form of Breathless (Btl-GFP; green) is also shown. The Rab11 compartments, which contain intraluminal vesicles (ILVs; green) and dense-core granules (DCGs; dark grey), and the late endosomes and lysosomes (magenta) are marked. **(b)** Basal wide-field fluorescence and differential interference contrast (‘Merge’) views of living secondary cells (SCs) expressing GFP-tagged form of Breathless (Btl-GFP; green) with no other transgene (control); or also expressing the open reading frame of the human GAPDH protein (hGAPDH), an RNAi construct targeting *Drosophila* GAPDH1 (*gapdh1* – RNAi #1), or an RNAi construct targeting *Drosophila* GAPDH2 (*gapdh2* – RNAi #1). SC outline approximated by dashed white circles, and acidic compartments are marked by LysoTracker Red (magenta). Btl-GFP-positive intraluminal vesicles (ILVs; green in ‘Merge’; grey in ‘Zoom’) are apparent inside compartments, surrounding dense-core granules (DCGs; asterisk in ‘Zoom’) and connecting DCGs to the limiting membrane of the compartment (yellow arrowheads, except in *GAPDH2* knockdown, where ILVs only surround peripheral small DCGs). DCG compartment outline is approximated by white circles. **(c)** Confocal transverse images of fixed accessory gland (AG) lumens from the same genotypes, containing Btl-GFP fluorescent puncta (yellow arrows). **(d)** Bar chart showing percentage of DCG compartments per cell containing clustered Btl-GFP-positive ILVs that are not directly in contact with DCGs. **(e)** Bar chart showing the percentage of DCG compartments per cell containing single spherical DCG. **(f)** Bar chart showing number of Btl-GFP fluorescent puncta in the lumen of AGs for the different genotypes. All data are from six-day-old male flies shifted to 29°C at eclosion to induce expression of transgenes. Genotypes are: *w; P[w^+^, tub-GAL80^ts^]/+*; *dsx-GAL4, P[w^+^, UAS-btl-GFP]/+* with no expression of other transgenes (control), or with UAS-hGAPDH, UAS-*gapdh1*-RNAi or UAS-*gapdh2*-RNAi overexpression. Scale bars in **(b)** (5 µm), in ‘Zoom’ (1 µm), and in **(c)** (20 µm). *** p <0.001, ** p < 0.01 and * p < 0.05 relative to control, n=24-34 cells, (f) n=10 glands.

To test the role of GAPDH in normal exosome biogenesis, we knocked down the two *Drosophila GAPDH* genes in SCs. There was no significant effect of *GAPDH1* knockdown on ILV biogenesis, although secretion was reduced **(Fig. 2b, 2c & 2f)**.

However, *GAPDH2* knockdown led to a severe disruption of dense-core granule formation in Rab11 compartments. There was no central dense core, but multiple small dense-core granules formed near the limiting membrane (**Fig. 2b & 2e)**. Btl-GFP-labelled ILVs were only located in close proximity to the limiting membrane and around the small DCGs; they were not observed in DCG-independent clusters in the compartment lumen as in controls **(Fig. 2b & 2d)**. A similar phenotype was observed with CD63-GFP-marked ILVs, where the proportion of Rab11 compartments making ILVs was also significantly decreased (**Extended Figure 4e**). In addition, *GAPDH2* knockdown significantly reduced exosome secretion from Rab11 compartments into the AG lumen (**Fig. 2f, Extended Fig. 4f**). This phenotype was not recapitulated by knocking down other enzymes, namely *Phosphoglucomutase 2* (*Pgm2a*), *Phosphoglucose isomerase* (*Pgi*) and *Phosphofructokinase* (*Pfk*), in the glycolytic pathway, suggesting that the observed effects on EV biogenesis are not a consequence of general metabolic changes **(Extended Fig. 5)**. Overall our data indicate that the formation of ILVs and DCGs in Rab11 compartments of SCs may be functionally linked processes that are regulated by GAPDH2. Importantly, inhibition of GAPDH2 expression suppressed the generation of clustered ILVs and their secretion as exosomes, suggesting that this protein normally plays an essential role in exosome biogenesis and aggregation, consistent with the EV clustering phenotype observed upon human GAPDH overexpression.

In light of the physiological role of GAPDH in EV biogenesis, we reasoned that the binding of the GAPDH G58 peptide to the outer surface of EVs could be utilized as a unique, but generic, tool to attach therapeutic moieties to the surface of EVs. As a proof of principle study, we fused the G58 peptide to the double-stranded RNA binding domain (dsRBD) of TAR RNA binding protein 2 (TARBP2), which exhibits high affinity for short double-stranded RNAs such as siRNA^20^. The resulting fusion protein, designated as G58T, was expressed and purified from *E.coli* cells. Incubation of the protein with purified EVs resulted in high levels of protein binding to EVs (designated as G58T EVs). Gel shift assay and spectrofluorimetric analysis revealed highly efficient binding of the G58T EVs with siRNA (∼ 500-700 siRNA molecules per EV). Moreover, bound siRNA was protected from degradation by RNase A (**Extended Fig. 6a-c**). Confocal microscopy of N2a cells treated with fluorescently labelled G58T EVs bound Cy-3 siRNA revealed efficient uptake of complexes by the cells (**Fig. 3a)**. Gene silencing assays using GAPDH predesigned siRNA, however, revealed low levels of gene silencing (∼ 15%) in N2a cells treated with G58T EV siRNA (**Extended Fig. 6e**). Co-localization studies using lysotracker dye suggested entrapment of delivered siRNA in late endosomes (**Extended fig. 6d**). To overcome endosomal entrapment, we investigated the fusion of different endosomolytic peptides including TAT, HA2 and the arginine-rich peptide of flock house nodovirus (FHV) to the G58T protein. These peptides have previously been successfully used to enhance the release of endocytosed drugs from late-endosomes^21–23^. HA2 fusion protein could not be expressed due to toxicity of the peptide. Attachment of either two TAT peptides or FHV peptide to the G58T protein (designated as G58T(tat)2 and G58TF respectively) resulted in significantly improved activity with ∼35% and 60% silencing of endogenous genes (GAPDH and HTT) genes in the cells respectively **(Fig. 3b; Extended Fig. 6g)**. Further treatment of cells with chloroquine, an endosomolytic molecule^24^, enhanced gene silencing efficiency to around 80%, consistent with the release of entrapped siRNA from late endosomes **(Fig. 3c; Extended Fig. 6f & 6h)**. Taken together, these results show highly efficient loading and delivery of siRNA into cells by G58T EVs, resulting in potent gene silencing when combined with fusion of endosomolytic peptides to G58T protein.

**Figure 3.**
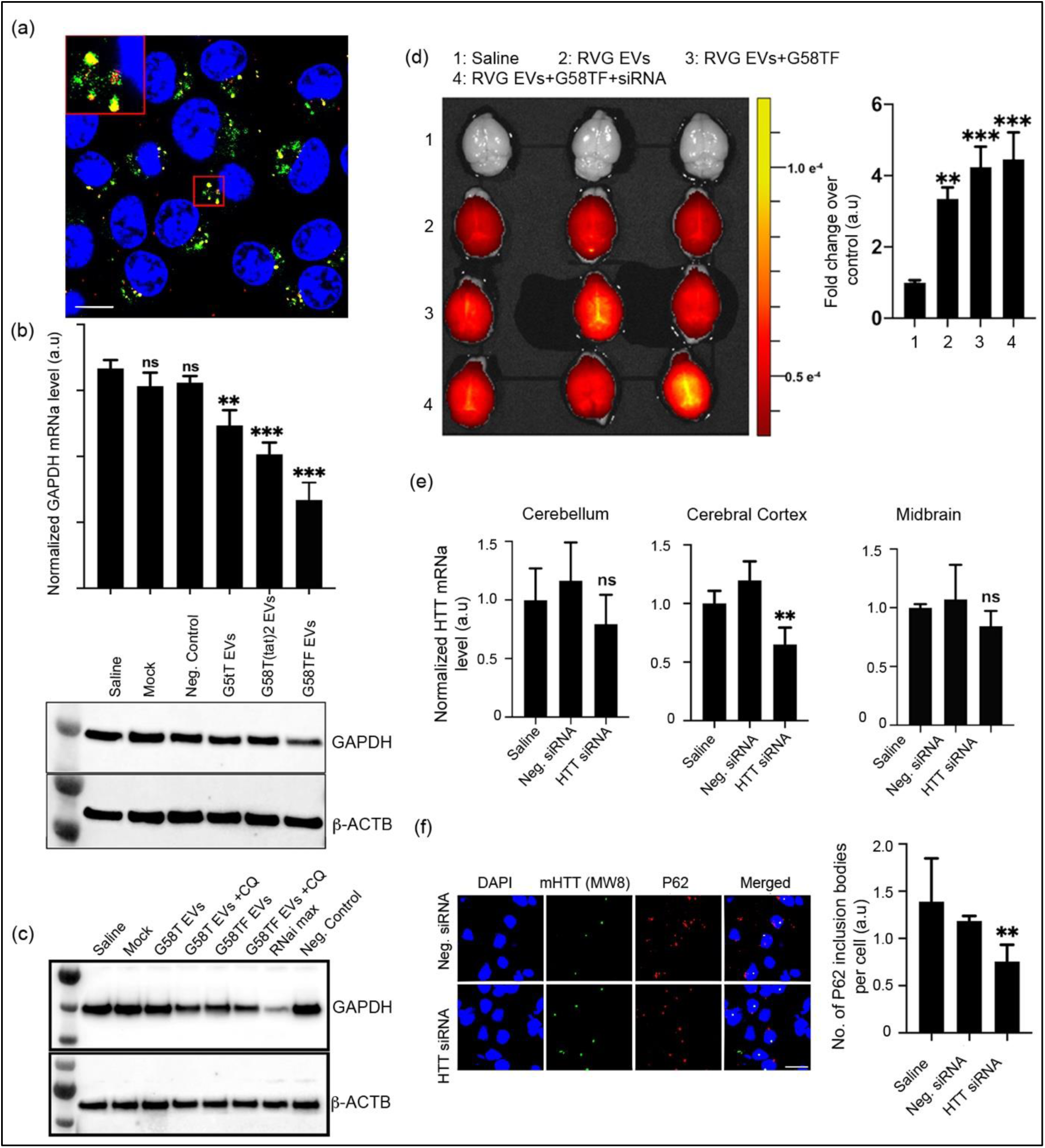
G58 peptide promotes EV-mediated siRNA delivery to the brain: **(a)** Confocal microscopy image of Neuro-2a cells after 4h of incubation with siRNA-loaded G58T EVs. Nuclei of cells were stained with Hoechst 3342. EV surface proteins were labeled with Alexa fluor-633 (red color) and siRNA was labeled with Cy-3 dye (green). Inset is the magnified image of the marked region showing colocalization of EVs with siRNA (yellow spots). Scale bar, 10µm. Representative image (n>3). Raw image is in Supporting data, Fig. 4. **(b)** Silencing of GAPDH expression in N2a cells by EVs engineered with G58T, G58T(tat)2 and G58TF proteins, respectively. 30nM of GAPDH siRNA was loaded on to EVs, which were added to cells. Top histogram represents GAPDH mRNA level determined after 48h of treatment, using probe-based qRT-PCR. Data were normalized with 18S rRNA relative to *GAPDH* mRNA in saline-treated cells. Mock: G58TF-bound EVs alone, Neg. Control: G58TF EVs + non-targeting siRNA. Results shown as mean ± s.d. **p=0.0006 and ***p<0.0001 when compared to control (one-way ANOVA; *n*= 3). Bottom western blot shows GAPDH protein level in N2a cells after 72 h of treatment. A single label is used for histogram and western blot images. Representative blot (n=3) **(c)** Western blot showing effect of chloroquine (CQ) on silencing of GAPDH protein by G58T and G58TF EVs in N2a cells. (Mock: G58TF EVs alone, Neg. Control: G58TF EVs + non-targeting siRNA, RNai max: siRNA with lipofectamine RNAiMAX reagent (Invitrogen)). 30µM chloroquine was added to cells along with the EVs. Representative blot (n =2) **(d)** *In vivo* fluorescence images of C57BL/6 mouse brains showing biodistribution of RVG-EVs, G58TF-RVG-EVs and G58T/siRNA-RVG-EVs. Images were taken after 4h of systemic administration of EVs. Surface proteins of RVG-EVs were labeled with cy5.5-NHS fluorescent dye. Mice injected with saline were used as a negative control. Histogram at the right side shows quantification of the fluorescent signal from the brains of treated mice. Results shown as mean ± s.d. **p=0.001 and ***p <0.0001 compared to control (saline) group (one-way ANOVA; n=3). **(e)** *In vivo* silencing of HTT gene in Q140 HD mouse model after systemic administration of G58TF EVs. Animals received four injections of EVs over four weeks. 72 h after the last dose, animals were sacrificed and sections of brain were analyzed for HTT mRNA level, using probe-based qRT-PCR. Data were normalized with 18S rRNA. Animals receiving saline were used to normalise HTT level in Neg. siRNA group (animals that received G58TF EVs + neg. siRNA) and HTT siRNA group (animals that received G58TF+HTT siRNA). Results shown as mean ± s.d. ns: non-significant, **p= 0.0012 when compared to control group. (one-way ANOVA; n =6) **(f)** Immunohistochemistry of cortical regions from the sacrificed animals. Images show mutant HTT protein level and p62-labeled inclusion bodies in the neurons of the cortex of HTT siRNA- and Neg. siRNA-treated animals. Histogram at the right shows quantification of p62 inclusion bodies in different groups. Results are mean ± s.d, n=8 slides chosen randomly from each group. **p=0.003 when compared to neg. siRNA group (two-way ANOVA)

To further assess the therapeutic potential of G58 engineered EVs in animal models, we investigated targeted delivery of siRNA into the mouse brain by co-expressing the RVG peptide on the EV surface. This peptide, which binds specifically to acetylcholine receptors, has been extensively used to deliver drugs into the brain^25^. Previous studies from our laboratory have shown that RVG increased brain accumulation of EVs without inducing toxicities in mice^26,27^. To determine whether binding of G58TF protein to the EV surface alters biodistribution of RVG-EVs, we assessed biodistribution of systemically administered RVG-EVs in C57 BL/6 mice. After 4h of administration, a significant fraction of drug dose of RVG-EVs was observed in the whole brains of mice. Binding of either G58TF or G58TF/siRNA to RVG-EVs did not change their biodistribution (**Fig. 3d)**, although, as anticipated, the majority of EVs were distributed to peripheral tissues, indicating rapid clearance of EVs from the blood **(Extended Fig. 7)**.

To assess the silencing efficiency and therapeutic potential of G58TF/siRNA RVG-EVs in the brain, we targeted the huntingtin gene in the Huntington’s disease (HD) mouse model Q140. HD is a neurodegenerative disease caused by mutation in the huntingtin gene that results in polyglutamine (poly Q) expansion in the protein, causing death of neurons^28^. Availability of HD mouse models provides a unique opportunity to assess therapeutic efficiency of siRNA-based drugs in silencing the mutant *HTT* gene^29^. Using a mixture of siRNAs targeting exon 1 of the HTT gene of Q140 mice, we systemically administered a total of four repeated EV doses weekly over 4 weeks. Analysis of brain tissues revealed almost 40% silencing of the HTT gene in mouse brain cortex and a highly significant decrease of p62 inclusion bodies in the cortical neurons of the treated animals, an important phenotypic HD marker **(Fig. 3f & 3g)**. p62 is an important regulatory protein of selective autophagy, and reduction in p62 aggregates in HD mouse models has previously been shown to restore HD-associated phenotypes^30^.

Taken together, our studies demonstrate a novel role for GAPDH protein in EV biogenesis and secretion of EVs from endosomal compartments, which appears to be associated with corresponding loading of GAPDH on EVs that can induce clustering. More importantly, by engineering GAPDH fusion proteins that have retained the G58 EV binding domain, we have developed a simple and highly robust method for therapeutic loading of RNA-based drugs on EVs for targeted drug delivery. These EVs can potentially be generated from any cell type. The method could be extended to deliver other types of cargoes, ranging from different types of nucleic acids including guide RNAs, phosphorodiamidate morpholino oligomers (PMOs), and plasmids to different types of proteins and peptides allowing the therapeutic potential of EVs to be enhanced and extended.

## Methods

### Isolation, purification and characterization of extracellular vesicles

#### (a) Isolation of EVs from cell culture media

Mesenchyme stem cells (MSCs), human embryonic kidney cells (HEK293T), human ovarian cancer cells (SKOVE-3), B-lymphoma cells (B16-F10) and HeLa cells were used to isolate extracellular vesicles. Cells were grown in DMEM GlutaMax medium (ThermoFisher Scientific, UK), containing 10% fetal bovine serum and 1X penicillin-streptomycin (PS, 100X contains 5000 U/ml of penicillin and 5000 µg/ml streptomycin; Sigma). MSCs were grown in RPMI GlutaMax media (ThermoFisher Scientific) with 10% FBS and 1X PS (Sigma). Cells were kept in an incubator maintained at 37°C with 5% CO_2_. For isolation of EVs, cells were seeded in 150 × 20 mm dishes (Star Labs, UK) at 10^6^ cells per dish in DMEM+10% FBS media. After 24 h of seeding, media of the cells was replaced with Opti-MEM reduced serum media (ThermoFisher Scientific, UK). Cells were incubated further for 48 h followed by collection of media in 50 ml falcon tubes (Sigma). To remove dead and floating cells, media was centrifuged at 500×g for 5 min. Supernatant was gently transferred into fresh tubes and centrifuged at 3000×g for 20 min at 4°C to pellet down cell fragments and remaining cell debris. Purified media was concentrated by tangential ultrafiltration (TFF), using 100 kDa Vivaflow 50R cartridges (Sartorius UK limited). Concentrated media was centrifuged at 10,000×g for 30 min to remove aggregates and bigger particles. The volume of media was further reduced to 2mL, using Amicon ultra-15 centrifugal filter units of 100 kDa MWCO (Millipore). Finally, the media was passed through a Sepharose 4 fast flow gel filtration column (GE Healthcare, 170149001), using the AKTA pure chromatography system (GE Health care). Purified EVs eluted from the column were used for the various biological assays. Generally, 500 ml of cell culture media were used at the beginning of the process and concentrated to 2ml before loading to the column.

#### (b) Isolation of EVs from hollow fibre bioreactor

For animal experiments, large quantities of EVs were isolated using hollow fibre bioreactor (FibreCell^®^ System, UK). 5 × 10^8^ MSCs and HEK293T cells were seeded into the extra-capillary space (ECS) of a hollow fibre bioreactor cartridge as per the guidelines of the manufacturer. Glucose level in the media was measured on a daily basis to monitor growth of cells. After one week of growth, extracellular material was harvested by gently flushing 25 ml of Opti-MEM media into ECS from the left end port to collect extracellular material on right end port of the cartridge. Withdrawn medium was flushed back and forth among the syringes, through the ECS, four times to dislodge material collected within the fibre bundles. The resulting harvest was used for isolation of EVs using differential centrifugation, ultrafiltration and size-exclusion chromatography as described above.

### Biophysical characterization of extracellular vesicles

#### (a) Nanosight Tracking Analysis

To measure size distribution and number of extracellular vesicles, nanoparticle tracking analysis was performed with the NS500 nanoparticles analyser (NanoSight, Malvern, Worchestershire, UK), as described previously^31^. Different dilutions of EVs ranging from 1:500 to 1:1000 in PBS were used to achieve a particle count of between 2× 10^8^ to 2× 10^9^ ml^−1^. During the recordings, a camera level of 12-14 was used. Using a script control function, three 30 s videos of each sample were recorded. After each video, a fresh sample was injected into the stage followed by a delay of 7 s to reduce turbulence of the flow. NTA2.3 software provided by the manufacturer was used to analyse and measure size and concentration of the EVs. All NTA measurements were carried out in triplicate. Data were analysed using GraphPad Prism 8 software.

#### (b) Electron microscopy of extracellular vesicles

For electron microscopy of extracellular vesicles, samples were prepared as per the protocol described previously^32^. Briefly 20 µl drops of purified EVs were added to parafilm and formvar-carbon coated EM grids (Agar Scientific, Stansted, UK) were floated on the top of EV droplets for 10 min with the coated side of the grid facing the drop. After the incubation, the grids were carefully transferred to PBS droplets for washing. Finally, the grids were placed briefly on water droplets to remove excess of salts. For contrasting, grids were initially transferred to a 50µl drop of uranyl-oxalate solution, pH 7, for 5 min, followed by embedding in 2% uranyl acetate for 5 min. Next, the grids were carefully removed and excess of fluid was removed by gently pushing the grid sideways on Whatman no. 1 filter paper. The grids were air dried and visualized under a JEOL 1010 transmission electron microscope at 120kV on EFI Tecnai 12 TEM (JEOL, Peabody, MA, USA).

#### (c) Western blotting of extracellular vesicles

For western blotting, EV lysates corresponding to 5–10 µg of total protein were run into 4-12% Bis-Tris plus gels (Invitrogen, ThermoFisher Scientific). Prior to loading, EV proteins were denatured by adding NuPAGE™ LDS Sample loading dye and reducing agent (ThermoFisher Scientific) to the EVs and heated at 70°C for 10 min. Samples loaded on the gel were run at 150V for 1 h in ice-cold NuPAGE™ MES running buffer. Proteins resolved on the gel were transferred onto 0.45 µm Immobilon-P, PVDF Membrane (Millipore), using a mini transfer blot cell (Bio-Rad). Membranes were blocked with SuperBlock™ T20 (TBS) Blocking Buffer (ThermoFisher Scientific) for 1 h at room temperature with gentle shaking. After blocking, primary antibodies diluted in the blocking buffer were added to the membrane and incubated overnight at 4°C on a shaker. Details of the antibodies used during the western blotting of EVs are given in the table below. After the incubation, the membranes were washed 3 times for 5 minutes each, using 1X tris-buffered saline, pH 7.6 (TBST, 2.4 g Tris base, 8.8 g of NaCl and 0.1% Tween). After the final wash, membranes were incubated with horseradish peroxidase (HRP)-conjugated secondary antibodies for 1 h at room temperature with gentle shaking. Post-incubation, membranes were washed 3 times for 5 min each with 1X TBST buffer and developed by chemiluminescence (GE healthcare, RPN2106). The Odyssey FC imaging system (LI-COR) was used to visualize the bands on the membrane. Image studio software provided with the system was used to analyse the data.

For western blotting of proteins from cells, radioimmunoprecipitation assay (RIPA) buffer (ThermoFisher Scientific) containing 1X protease inhibitor (Halt protease inhibitor, ThermoFisher Scientific) was added to cells and incubated on ice for 10 min. Lysates of the cells were passed through 26G needle syringes several times and centrifuged at 12,000×g for 20min to remove genomic DNA aggregates and insoluble lipids. Supernatant of the cell lysate corresponding to 5-10 µg of the total proteins were used for the western blotting.

| **Antibodies** | **Source** | **Dilution** |
| --- | --- | --- |
| Alix | Santa Cruz Biotechnology, sc-53540 | 1:500 |
| CD81 | Santa Cruz Biotechnology, sc-166029 | 1:500 |
| GAPDH | Sigma, G9545-100µl | 1:1000 |
| TRBP | Abcam, ab72110 | 1:1000 |
| cMyc | Abcam, ab32 | 1:1000 |
| Flag | Sigma, F2555-100UL | 1:1000 |
| GFP | Abcam, ab38689 | 1:1000 |
| LAMP | Abcam, ab18529 | 1:1000 |
| Lactoferrin | Abcam, ab109216 | 1:1000 |
| Beta Actin | ThermoFisher Scientific, MA5-15739 | 1:2000 |
| Anti-Rabbit IgG-HRP | Sigma, A0545-1mL | 1:2000 |
| Anti-Mouse IgG-HRP | Sigma, A5278-1mL | 1:5000 |
| MMP-2(2c1) | Santa Cruz Biotechnology, sc-13594 | 1:500 |

### Purification of GAPDH and G58T fusion proteins from bacterial cells

#### (a) GAPDH protein

The cDNA sequence of human GAPDH (OriGene, UK) was inserted into the pET-28b(+) vector (Novagen). At the N-terminus of GAPDH, six residues of histidine were inserted for purification by Ni-NTA chromatography. At the C-terminus, a FLAG-tag was attached. The vector was transformed into BL21 (DE3) competent cells (New England Biolabs, UK) for expression. 1 mM Isopropyl β-D-1-thiogalactopyranoside (IPTG, Sigma) was used to induce expression of the gene. Cells were grown at 37°C during the expression. GAPDH protein was purified by Ni-NTA chromatography (Qiagen) under native conditions, using standard protocols recommended by the manufacturer. Ice-cold sodium-phosphate buffer, pH 8.0 (50mM Na_2_HPO_4_/NaH_2_PO_4_, 10mM Tris-Cl, 300mM NaCl, 5mM imidazole) was used during resuspension and lysis of the BL21(DE3) cells. Purified GAPDH protein was eluted with 250 mM imidazole in sodium phosphate buffer, pH 6.0. Protein was dialyzed and passed through a sephacryl s-200 HR column (GE Health Care) for further purification. The protein was analysed by SDS-PAGE. Size of the protein was confirmed by mass spectrometry using MALDI.

#### (b) G58T, G58T(tat)2 and G58TF proteins

The DNA sequence encoding theG58 peptide (amino acid residues 58 to 100 of human GAPDH) was fused with second domain of the human TAR RNA binding protein (TRBP2, domain sequence was from UniProtKB; Sequence ID: Q15633). The DNA sequence of the fused G58 and TRBP2 second domain was synthesized commercially, using gBlcoks^®^ Gene Fragments tools (Integrated DNA technology, Belgium). The DNA sequence of TAT and arginine-rich peptide of flock house virus alpha capsid protein was custom-synthesized (integrated DNA technology, Belgium) and fused to the TRBP2 second domain by PCR cloning. The arrangement of domains in different G58 fusion proteins is as follows:

G58T: His_6_…G58 peptide….(Gly_4_Ser)….TRBP2 second domain

G58T(tat)2: His_6_…G58 peptide….(Gly_4_Ser)….TRBP2 second domain…(PKR linker).. (TAT) … (TAT)**^2^**

G58TF: His_6_…G58 peptide….(Gly_4_Ser)….TRBP2 second domain…(PKR linker)**^1^**..(FHVpeptide)**^3^**

1. PKR linker: Amino acid sequence between first and second RNA-binding domain of human protein kinase R
2. TAT peptide sequence: GRKKRRQRRRPQ
3. FHV peptide sequence: GRRRRNRTRRNRRRVRG^33^

The DNA sequences of these fusion proteins were cloned into the pET-28b(+) vector. G58T was expressed in BL21 (DE3) cells, using similar conditions to those mentioned above. G58T(TAT)_2_ and G58TF were expressed in BL21 (DE3) Rosetta (Novagen, Merk, USA).

The cells were grown normally at 37°C until OD_600_ reached between 0.5 to 0.6. At this point, 0.2mM IPTG was added to the culture, and bacteria were grown at 21°C for 10 to 12 h. All versions of G58 fusion proteins were purified by Ni-NTA chromatography under denaturing conditions, using 6M urea. Purified proteins were analysed by SDS PAGE and MALDI. Proteins were refolded by the dilution method. Briefly, proteins bound to Ni-NTA resin were eluted by 250mM imidazole in sodium phosphate buffer, containing 2M urea. Eluted proteins were further diluted with an equal volume of ice-cold PBS containing 5% glycerol and 0.1 mM DTT. After 5 min of incubation, the protein was added dropwise to PBS (with constant stirring) to reduce the urea concentration to 0.5 M. After the dilution, aggregated proteins were removed by centrifugation at 30,000×g for 20 min (Beckman Coulter, USA). Protein was concentrated by centrifugal spin filters (Millipore) and passed through a PD10 (GE Healthcare, USA) column to remove final traces of urea.

### Binding of GAPDH and G58T fusion proteins to extracellular vesicles

For exogenous binding of GAPDH to EVs, purified EVs ranging from 7.5 ×10^11^ particles/ml to 1×10^12^ particles/ml were used for incubation with GAPDH and G58 fusion proteins. Proteins in the range of 5 to 20nmoles were added to each pmoles of EVs and incubated for 2h at 4°C on a rotor top. Unbound proteins were removed by gel filtration. Proteins were also added directly to concentrated cell culture media and mice serum for binding followed by gel filtration to remove the unbound proteins from the EVs. Binding of GAPDH proteins to EV surface was monitored by UV-280 chromatogram (Akta pure, GE Healthcare), western blotting, GAPDH kinetic assay and gel-shift assay.

### GAPDH kinetic assay

A fluorescence-based KDalert^TM^ GAPDH assay (Invitrogen, Life Technologies) was used to measure enzymatic activity of GAPDH protein of extracellular vesicles. The assay measures the conversion of NAD^+^ to NADH in the presence of phosphate and glyceraldehyde-3-phosphate (G-3-P). Under the recommended assay conditions, the rate of NADH production is proportional to the amount of GAPDH enzyme present. A given number of EVs were added to the substrate in a 96-well plate. A fluorescent microplate reader (Clariostar Plus, BMG Labtech) was used to acquire data under kinetic mode, using λ_ex_ 560nm and λ_em_ 590nm. Data were analyzed using software (MARS Data Analysis) provided with the microplate reader.

### Gel shift assay and RNase A protection assay

Binding of G58T, G58T(TAT)2 and G58TF EVs to siRNA was assessed by gel shift assay. 20 pmole of siRNA was added to a given number of EVs in PBS buffer and incubated for 10 min at room temperature. After the incubation, complexes were loaded onto a 2% agarose gel stained with 0.5 µg/ml ethidium bromide for visualization under a UV-illuminator. Binding of siRNA to EVs was determined by analyzing the shift of siRNA on the gel. siRNA bound to EVs remained trapped near the wells while free siRNA migrated freely in the agarose gel. Using NTA data and the gel binding assay, number of siRNAs bound to EVs was determined.

For the RNase A protection assay, 0.2mg/mL of RNase A (Qiagen) was added to EV-siRNA complexes and incubated for 6 h at 37°C. The enzyme was inactivated by adding an equal volume of hot SDS lysing solution (2% SDS, 16 mM EDTA) and incubated at 100°C for 5 min. siRNA was isolated using TRIzol reagent (Invitrogen, Life Technologies) and analysed by agarose gel electrophoresis.

### Confocal Imaging

To determine uptake of siRNA-loaded EVs into cells, N2a cells were seeded onto coverslips in 6-well tissue culture plates (Sigma). Surface proteins of EVs were labelled with Alexa Fluor 633 (Invitrogen, Life Technologies). siRNA^Cy3^ was custom synthesized (Integrated DNA technologies, Belgium). After 24 h of seeding cells, complexes of Alexa Fluor 633-labelled EVs and siRNA^Cy3^ (20 pmoles) were added to the cells and incubated for 6h. After the incubation, cells were washed with PBS containing 20U/ml heparin sulphate (Sigma), stained with Hoechst 33258 (Molecular probes, Life technologies) and fixed with 4% paraformaldehyde (ThermoFisher scientific, UK). The coverslips were mounted on glass slides using Ibidi mounting medium (Ibidi, Germany) and visualized under fluorescence laser confocal microscope, using a 60X oil immersion objective lens (FV 1000, Olympus). The data were assessed by using FV100 software supplied with the microscope.

### *In vitro* gene silencing by G58-modified extracellular vesicles

For silencing of genes in N2a cells and HeLa cells, G58T, G58T(TAT)2 and G58TF proteins were incubated with either MSC or HEK293T EVs. After binding, purified EVs were analyzed for siRNA binding, using gel shift assay, to determine the amount of EVs needed to completely bind a given amount of siRNA. Using that ratio, a given siRNA concentration bound to EVs was added to the cells in complete DMEM GlutaMax medium. Details of the siRNA used for gene silencing experiment are given in the table below. After 24 h of incubation, medium was changed and cells were further incubated in the medium. For mRNA analysis, cells were harvested after 48 h of treatment, using TRIzol (Invitrogen, Life Technologies). 250 ng of total RNA was used for reverse transcription PCR (RT-PCR), using the PrimeScript reverse transcriptase kit (Takara, Japan). Probe-based quantitative real time PCR was used to assess quantities of GAPDH and HTT mRNA. Amplification of 18S rRNA and Hypoxanthine-guanine phosphoribosyl transferase (HPRT) mRNA was used for internal controls. LinRegPCR software was used to determine the efficiency of PCR. Data were analyzed by PfafEl analysis, as described. GraphPad Prism 8 software was used to determine percentage of gene silencing and level of statistical significance among the treatment groups, using one-way ANOVA. Details of gene-specific probes are given in the table below.

For immunoblotting, cells were lysed after 72h of treatment with RIPA buffer, containing IX protease inhibitor cocktail (ThermoFisher Scientific). 5μg of total cell lysate proteins were resolved on 4-12% SDS-PAGE and transferred onto 0.45 μm polyvinyl difluoride membrane (Millipore) as described above under western blotting of EVs.

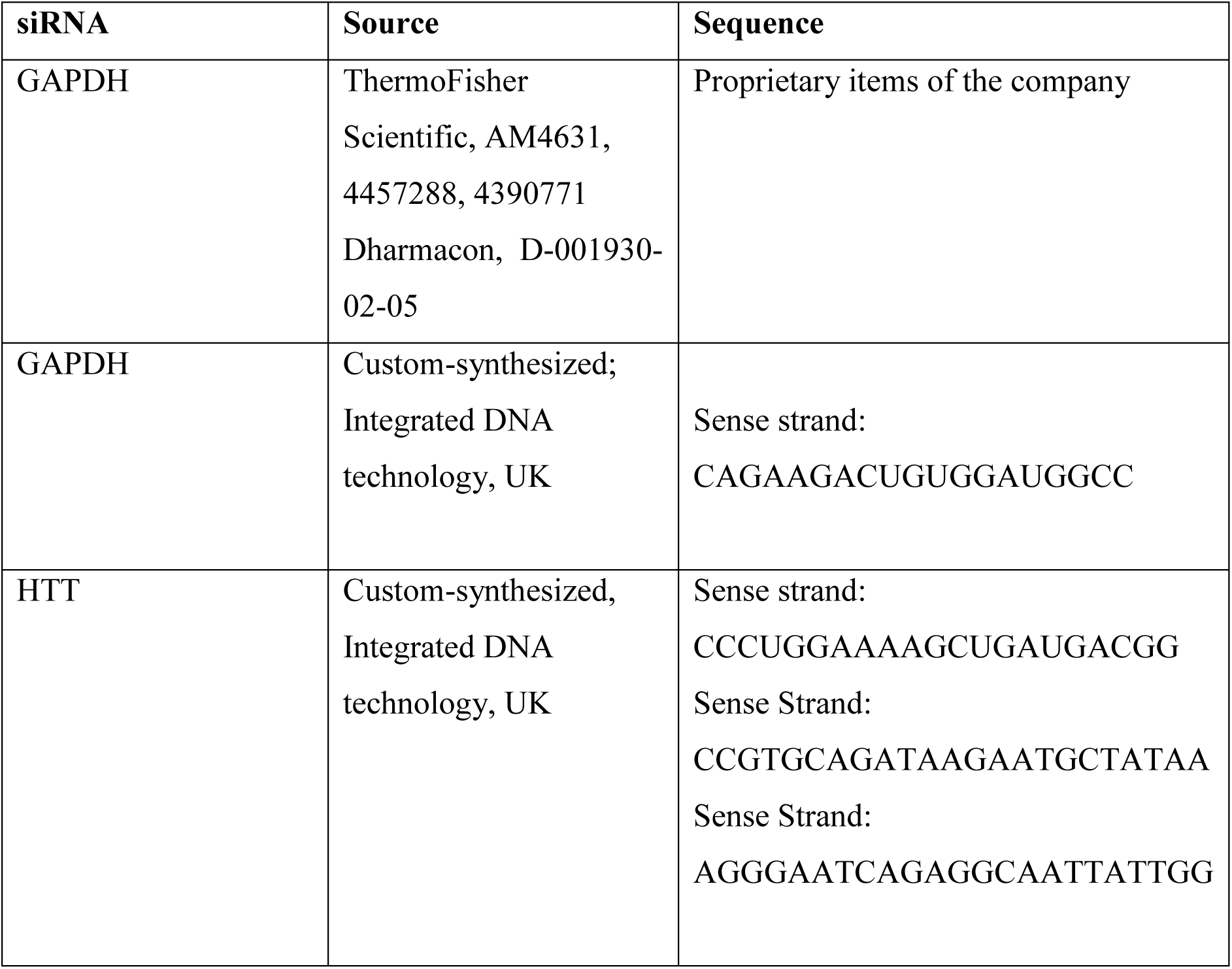
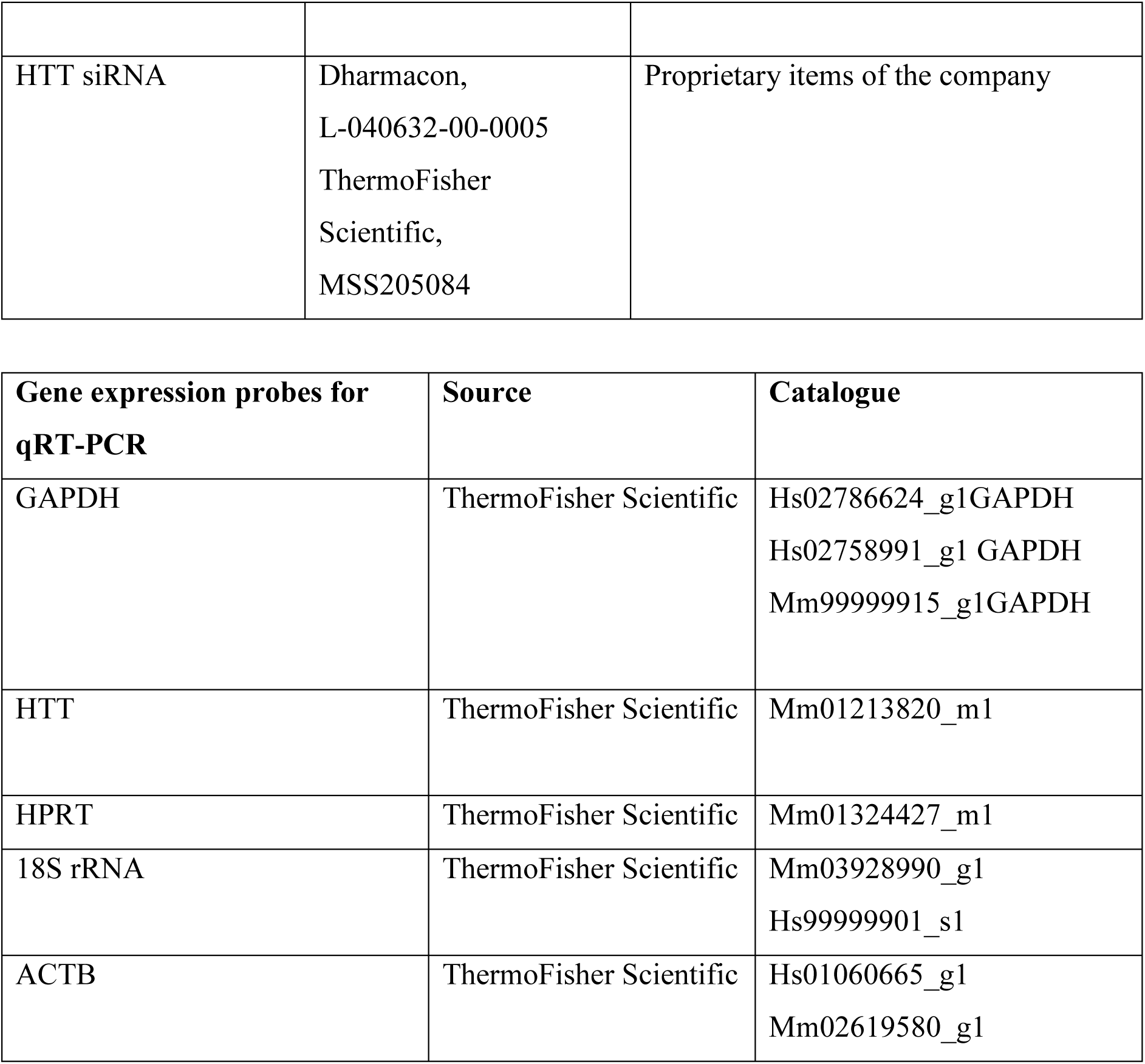

### Animal Experiments

#### (a) Biodistribution of Extracellular vesicles

Six-week-old female C57BL/6 mice were used. All animal experiments were done in compliance with the standards set down in the Animals (Scientific Procedures) Act 1986, revised 2012 (ASPA, UK). Mice were grouped randomly into 4 groups, each group containing three mice. HEK293T cells stably transfected with RVG-LAMP-2B protein were seeded into 15×1.2 cm tissue culture plates for isolation of EVs. Immunoblotting of EVs and HEK293T cell lysates was carried out to assess expression of RVG peptide on surface of EVs. RVG-expressing EVs were labeled with sulpho-Cy5.5 NHS ester dyes (Abcam, ab235032). Briefly, 2×10^12^ EVs in 1 ml PBS buffer were incubated with 100 µmoles of sulpho-Cy5.5 NHS ester, which was prepared in DMSO as a 10 mM stock. EVs were incubated at room temperature for 1 h with gentle shaking. After 1 h of incubation, G58TF proteins was added to EVs and incubated for 2 h at 4°C. Excess of dye and protein was separated by passing EVs through a gel-filtration column. After concentrating EVs, the following single doses of EVs were given into the animals. Doses were calculated based on previously reported EV biodistribution studies^27^.

| S/No | Composition of the formulation | Number of EVs /g body weight | Amount of siRNA |
| --- | --- | --- | --- |
| Group I | Saline | ———— | ———— |
| Group II | RVG EVs | $(1.5-2.0) \times 10^{10}$ | ———— |
| Group III | EVG EVs +G58TF | $(1.5-2.0) \times 10^{10}$ | ———— |
| Group IV | EVG EVs<br>+G58TF+siRNA | $(1.5-2.0) \times 10^{10}$ | 10µg (0.5mg/kg dose) |

After 4 h of administration, animals were sacrificed and organs harvested were visualized under an *in vivo* animal imaging system (IVIS, Perkin Elmer) for Cy5.5 fluorescence. The data were analyzed with the IVIS software (Living Image Software for IVIS)

#### (b) Silencing of huntingtin gene (HTT) gene in Q140 Huntington disease model animals

Huntington’s disease model Q140 mice of 1-year age were used to assess silencing of mutant and wild HTT gene silencing by intravenous administration of siRNA-loaded EVs. In the Q140 mouse model, exon 1 of mouse HTT is humanized and contains 140 CAG repeats. The mice have a slow progression of disease phenotype, which starts to appear at the age of 6 months. In this experiment, Q140 mice of 1-year age were grouped into saline, negative and treatment control groups. Each group contained 6 mice. Saline group was used as a control to assess levels of HTT mRNA after administering EVs loaded with either negative siRNA (Negative group) or a mixture of HTT siRNA (treatment group). RVG EVs bound to G58TF protein were used for HTT silencing experiment. EV doses were calculated based on 0.5 mg/kg siRNA dosage regimen. Number of EVs needed to bind a given amount of siRNA were calculated by gel-shift assay. 150-200 µl of EVs were administered intravenously. A total of 4 doses were given to animals. Animals received each dose weekly. 72h after the last dose, animals were euthanized and different sections of the brain were analyzed for HTT mRNA and protein expression. Levels of HTT mRNA in the saline treated group were assigned as 1 and based on that, the percentage of HTT silencing was determined by using LinRegPCR PfafE1 analysis. For determining the level of HTT protein in the brain tissues, Agarose gel electrophoresis for resolving aggregates (AGERA) was carried out to detect mutant HTT protein aggregates. However, we could not analyze the immunoblot due to high background. Immunohistochemistry of the cortex regions of the brain was carried out to determine the level of mutant HTT protein aggregates and p62 inclusion bodies. Data were analyzed by GraphPad Prism software. Multivariate ANOVA (2 tails) and post-hoc adjustment using Dunnett’s test was used to calculate statistical significance between the means of the groups.

### *Drosophila* Stocks and Genetics

The following fly stocks, acquired from the Bloomington *Drosophila* Stock Centre, unless otherwise stated, were used: *UAS-CD63-GFP*^34^ (a gift from S. Eaton, Max Planck Institute of Molecular Cell Biology and Genetics, Dresden, Germany), *dsx-GAL4*^35^ (a gift from S. Goodwin, University of Oxford, UK), *tubulin-GAL80^ts^*; *UAS-Btl-GFP*^36^, *UAS-gapdh1-RNAi* (#1, TRiP.GL01094; #2, TRiP.HMC05219)^37^; *UAS-gapdh2-RNAi* (#1, TRiP.JF02072; #2, v106562, Vienna *Drosophila* Resource Centre [VDRC]^38^), *UAS-Pgi-RNAi* (TRiP.HMC03362), *UAS-Pgm2a-RNAi* (TRiP.HMC04927), and *w^1118^* (a gift from L. Partridge, UCL, UK).

Flies were reared at 25°C in vials containing standard cornmeal agar medium (12.5 g agar, 75 g cornmeal, 93 g glucose, 31.5 g inactivated yeast, 8.6 g potassium sodium tartrate, 0.7 g calcium, and 2.5 g Nipagen (dissolved in 12 ml ethanol) per litre. They were transferred onto fresh food every two days. No additional dried yeast was added to the vials. Temperature-controlled, SC-specific expression of *UAS-CD63-GFP* and *UAS-Btl-GFP* was achieved by combining these transgenes with the specific driver, *dsx-GAL4* and the temperature-sensitive, ubiquitously expressed repressor *tubulin-GAL80^ts^*. Newly enclosed virgin adult males were transferred to 29°C to induce post-developmental SC-specific expression. For overexpression and knockdown experiments, the same strategy was employed, but in the presence of a UAS transgene. Six-day-old adult virgin males were used throughout this study to ensure that age- and mating-dependent effects on SC biology were mitigated^18,19,39^.

### Preparation of Accessory Glands for Live Imaging and Exosome Secretion Analysis

Adult male flies were anaesthetised using CO_2_. The abdomens were removed from anaesthetised flies and submerged in PBS (Gibco^TM^). The whole male reproductive tract was carefully pulled out of the body cavity. The accessory glands were isolated by separation of the ejaculatory bulb from the external genitalia, fat tissues and gut. Finally, the testes were removed by scission close to the seminal vesicles, as they often folded over the accessory glands, obscuring imaging.

The isolated accessory glands were transferred to a 9-spot depression glass plate (Corning^®^ PYREX^®^) containing ice-cold PBS and kept on ice until sufficient numbers had been obtained. They were then stained with ice-cold 500 nM LysoTracker^®^ Red DND-99 (Invitrogen^®^) (1:1000) in 1 × PBS for 5 min. Finally, the glands were rinsed in ice-cold PBS before being mounted onto high precision microscope cover glasses. A custom-built holder secured the cover glasses in place during imaging by wide-field microscopy. To avoid dehydration and hypoxia, the glands were carefully maintained in a small drop of PBS, surrounded by 10S Voltalef^®^ (VWR Chemicals), an oxygen-diffusible hydrocarbon oil^40^, and kept stably in place by the application of a small cover glass (VWR).

### Imaging of *Drosophila* Secondary Cells (SCs)

#### a) Wide-Field Microscopy

For wide-field imaging, living SCs were imaged using a DeltaVision Elite wide-field fluorescence deconvolution microscope (GE Healthcare Life Sciences) equipped with a 100X, NA 1.4 UPlanSApo oil objective lens (Olympus), and a Cool SNAP HQ2 CCD camera (Photometrics^®^). The images acquired were typically z-stacks spanning 8-12 µm depth with a z-distance of 0.2 µm. Images were subsequently deconvolved using the Resolve 3D-constrained iterative deconvolution algorithm within the softWoRx 5.5 software package (GE Healthcare Life Sciences).

#### b) Confocal Microscopy

Confocal images of accessory glands were acquired by using LSM 880 laser scanning confocal microscope equipped with a 63×, NA 1.4 Plan APO oil DIC objective (Carl Zeiss). RI 1.514 objective immersion oil (Carl Zeiss) was employed.

### *Drosophila* Exosome Secretion Assay

Virgin six-day-old males of each genotype described previously were dissected in 4% paraformaldehyde (Sigma-Aldrich) dissolved in PBS. The glands were left in 4% paraformaldehyde for 15 minutes to preserve the luminal contents before being washed in PBT (PBS containing 0.3% Triton X-100, Sigma-Aldrich) for 5 min. Glands were rinsed with PBS and then mounted onto SuperFrost^®^ Plus glass slides (VWR), removing excess liquid using a Gilson pipette, and finally immersed in a drop of Vectashield with DAPI (Vector Laboratories) for imaging by confocal microscopy.

Exosome secretion was measured by sampling within the central third of each gland. Identical microscope settings and equipment were used throughout. At each sampling location, ten different z-planes, spaced by 1 μm and at a distance from any SCs, were analysed.

The automated analysis of exosome secretions by SCs was performed using ImageJ2^41^, distributed by Fiji. The ‘Noise>Despeckle’ function was first used to remove background noise, followed by conversion to a binary image. The number of fluorescent particles was assessed using the ‘Analyse Particles’ function. The average number of particles for all stacks was then quantified and compared to controls.

### Analysis of ILV Content in Dense-Core Granule Compartments

Living SCs from six-day-old males of each genotype described previously were imaged using identical settings by wide-field microscopy. The total number of dense-core granule (DCG) compartments that contained clustered fluorescently labelled puncta was counted in each cell, using a full z-stack of the epithelium. Three individual SCs were analysed from each of ten to fifteen glands.

### Dense-Core Granule Compartment Analysis

Dense-core granule (DCG) compartment numbers were manually quantified in ImageJ2 by using complete z-stacks of individual cells acquired with differential interference contrast (DIC) microscopy on the wide-field microscope. The morphology and position of DCGs were also analysed, and each compartment was categorised by the absence or presence of a single spherical DCG. Three individual SCs per gland from each of ten to fifteen glands were used.

### Data Analysis

All data are representative from at least two to three independent experiments. The distribution of residuals was tested for normality using Q-Q plots and the appropriate statistical test was applied. For quantification of western blots, Image Studio Lite (Li-Cor) software was used. Adobe Photoshop CS4 software was used to crop and arrange the western blotting, confocal microscopy and electron microscopy figures. For quantitative real-time PCR, Linear regression (LinReg) PCR software was used to determine PCR efficiency using Microsoft excels. Nonparametric one-way ANOVA, using Bartlett’s and Brown-Forsythe test, was used to calculate F and P value. For mouse experiments, data were plotted and analysed, using multivariant (two-tail) ANOVA. For SC analysis and exosome biogenesis, the Kruskal-Wallis test was employed followed by Dunn’s test to compare individual control and experimental datasets. All data analyses and statistics were conducted using GraphPad Prism v8.0 (GraphPad Software Inc., La Jolla, CA, USA).

## Acknowledgements

We are grateful to I. Dobbie for technical expertise and support with microscopy, which was undertaken in the Wellcome Trust-funded MICRON Oxford Advanced Bioimaging Unit. We thank Errin Johnson for providing technical assistance in electron microscopy of extracellular vesicles at Electron microscopy facility, Sir William Dunn School of Pathology, University of Oxford. We also thank S. Eaton, S. Goodwin, as well as the Bloomington and Vienna Stock Centres for *Drosophila* stocks. This project received funding from Medical Research Council UK (MRC reference number: MR/M007715/1). For the *Drosophila* work, we acknowledge the support of the BBSRC (BB/K017462/1, BB/N016300/1, BB/R004862/1), Cancer Research UK (C19591/A19076), and the Wellcome Trust (Strategic Awards 102347/Z/13/Z).

## Author Contributions

G. H. Dar conceived of the presented idea, planned and carried out the experiments. C. Wilson, C. C. Mendes and G. H. Dar planned the *Drosophila* experiments. C. C. Mendes carried out and analysed the experiments in *Drosophila*. G. H. Dar and W. L Kuan designed and carried out HTT gene silencing experiments in Q140 mice. M. Conceição and G. H. Dar carried out the EV biodistribution experiments in C57BL/6 mice. G. H. Dar, I. Uliyakina and M. Lobo designed and carried out the immunoprecipitation experiments. T. Roberts, I. Mager and G.H. Dar analysed the animal data. G. H. Dar, C. Wilson and M. Wood wrote the manuscript. M. Wood supervised the study. All authors discussed the results and assisted in the preparation of the manuscript.

## Extended Figures

**Extended Figure 1.**
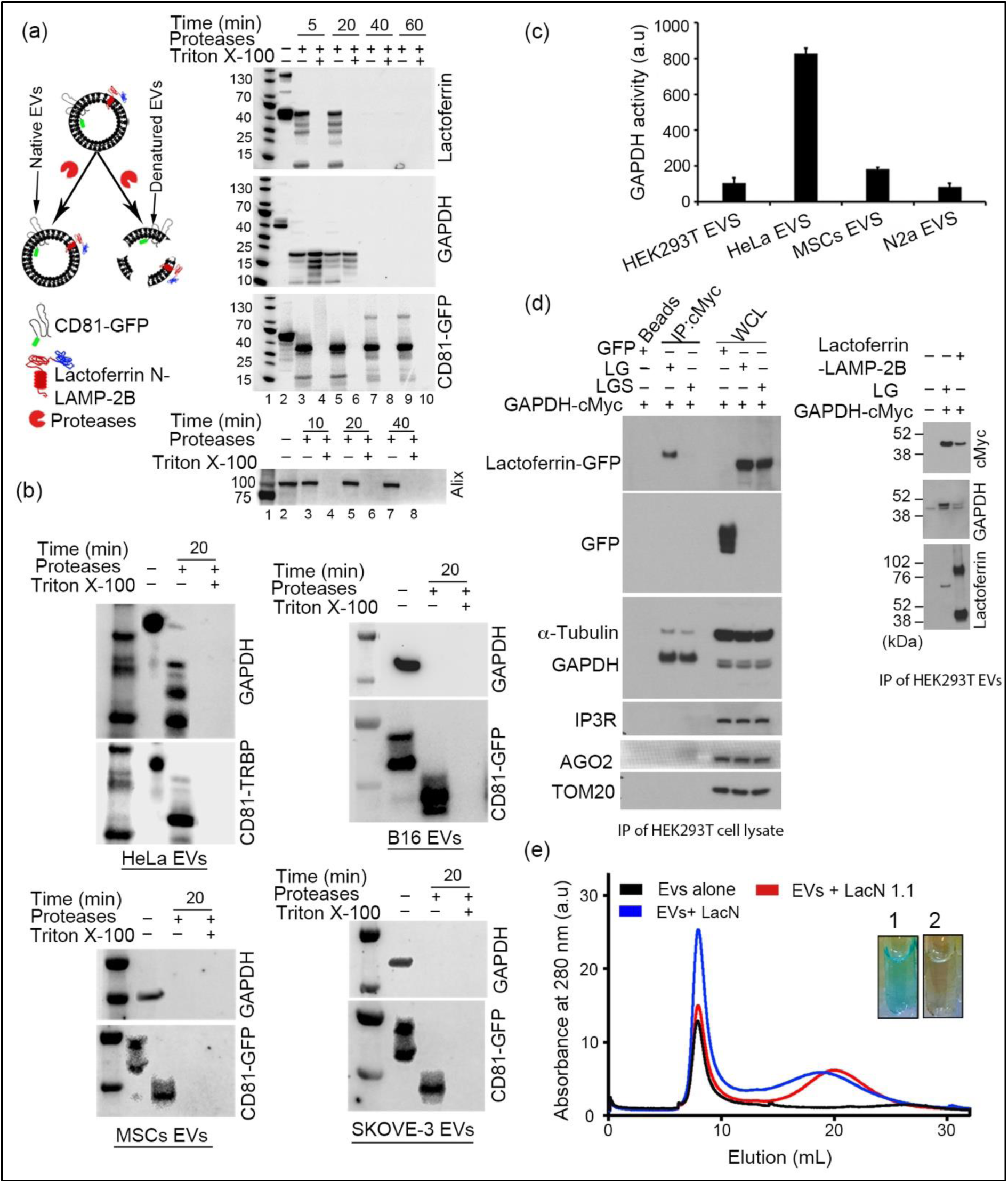
GAPDH present on outer surface of EVs binds to cleaved lactoferrin: **(a)** Protease digestion assay of EVs under native conditions and following detergent treatment (denatured conditions). Top left represents schematic drawing of protease digestion assay. CD81-GFP was used as positive control for protein extending into the lumen of EVs. After protease treatment, protein lysates were analysed with lactoferrin, GFP, GAPDH and Alix antibodies on western blots. Lane 1, standard protein marker; 2, non-treated EVs; 3, 5, 7 and 9, EVs incubated with pronase (mixture of proteases) for indicated times under native conditions; 4, 6, 8 and 10, EVs were treated with detergent and heated at 90 °C for 5 min to disrupt membranes followed by treatment with proteases. Assay was carried out at 37°C. Digestion of lactoferrin and GAPDH proteins under native conditions confirms their presence on outer surface of EVs. Presence of intact GFP-containing band, which is partly truncated due to cleavage of an extracellular CD81 domain, after incubating EVs for 60 min with protease under native conditions, confirms integrity of the membrane during the treatment. Bottom immunoblot shows the digestion pattern of Alix protein, which is an EV protein marker present in the lumen of the EVs. Under native conditions the protein remained intact, further confirming integrity of the EV membrane during the protease digestion assay. Representative image (n=3). **(b)** Western blots of EVs from different cells after protease digestion assay. Digestion of GAPDH protein under native and denatured condition confirms presence of GAPDH on the outer surface of EVs. CD81-GFP and CD81-TRBP were used as positive control for proteins extending into the EV lumen. Representative image (n=2). **(c)** GAPDH enzyme kinetics of EVs from different cell sources. Equal numbers of EVs were used to monitor in real time the formation of NADH by using the KDalert GAPDH assay kit (Invitrogen). Values are the fluorescence intensity of NADH measured under kinetic mode using 560 nm excitation and 590 nm emission. Data were normalized with the blank sample, containing substrate only. Results shown as mean ± s.d. (n=3**) (d)** Co-immunoprecipitation of lactoferrin and GAPDH protein expressed in HEK293T cells and EVs. Beads conjugated to cMyc antibody were used to capture GAPDH-cMyc protein in the cell lysate that was analyzed by western blotting, using lactoferrin antibody. Detection of lactoferrin-GFP band by the GFP antibody in lane 2 (top upper blot) confirms interaction of GAPDH protein with lactoferrin N domain. LG, lactoferrin-GFP protein; LGS, Lactoferrin-GFP with signal peptide of LAMP-2B. Attachment of a signal peptide to LG protein resulted in loss of lactoferrin interaction with the GAPDH for reasons that were not investigated; WCL, crude cell lysate used as positive control. Presence of α-tubulin bands shows interaction of GAPDH with α-tubulin. Insulin triphosphate receptor 3 (IPR3), Argonuate 2 (AGO2) and mitochondrial import receptor subunit TOM20 were used as negative controls. Western blot image on the right side shows co-immunoprecipitation of lactoferrin and GAPDH on surface of EVs. cMyc beads were used to capture GAPDH-cMyc fusion protein from EVs. Captured proteins were analyzed by western blotting using GFP, lactoferrin and GAPDH antibodies. Western blotting image shows presence of lactoferrin-GFP and lactoferrin-LAMP-2B bands, confirming their interaction with GAPDH proteins. Representative image (n=3). **(e)** Chromatogram of gel filtration chromatography showing absorbance of EVs at 280 nm after incubation with Lactoferrin N1.1 (lacN1.1) and Lactoferrin N (LacN) respectively. Increase in absorbance (OD280) of EVs bound to LacN is due to binding of lactoferrin N to surface of EVs. Inset represents EVs bound to Alexa Fluor-633 labelled lactoferrin N protein (designated as 1). Tube labeled with 2 represents EVs alone. Representative image (n>3).

**Extended Fig. 2.**
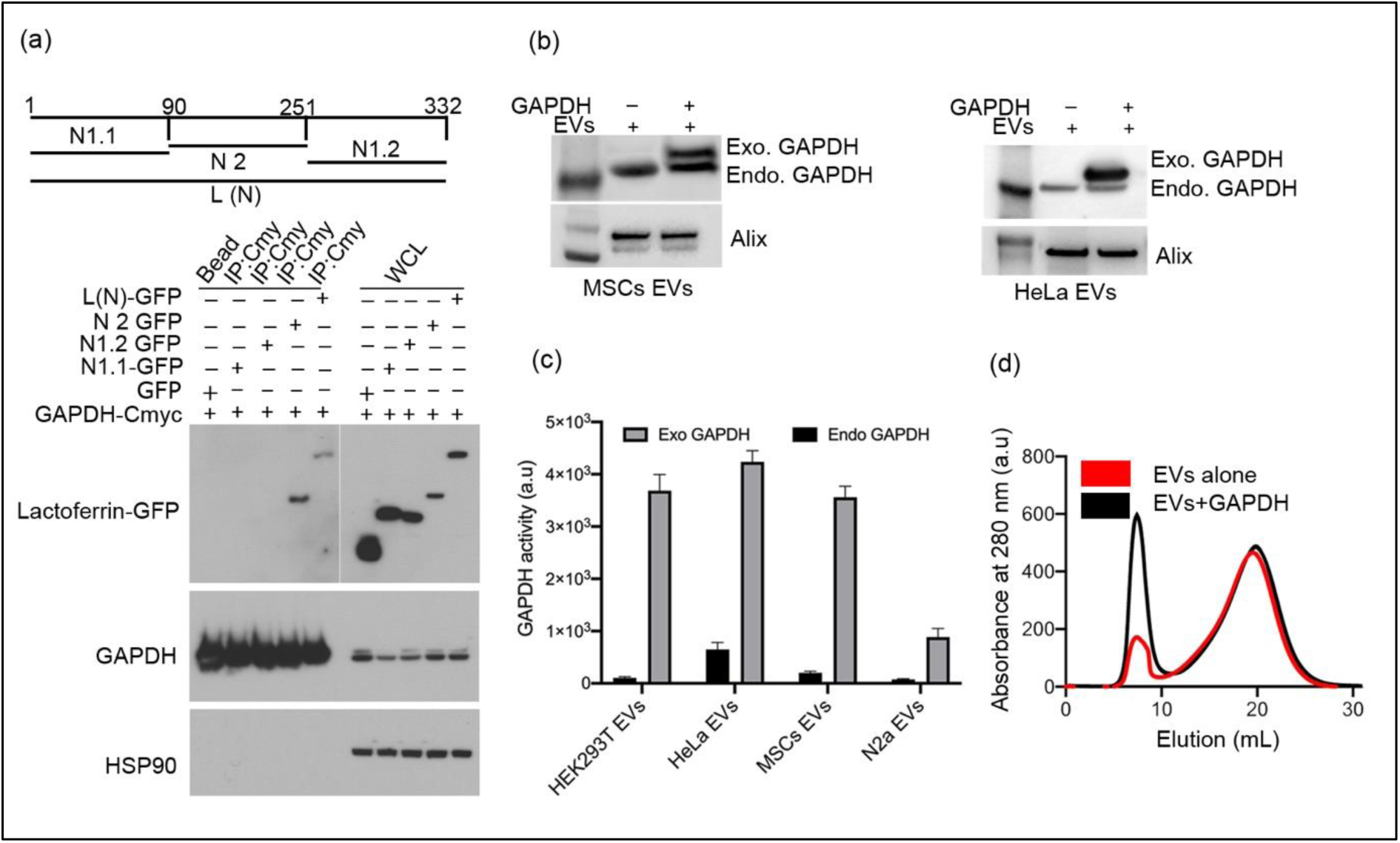
Binding of Exogenous GAPDH to EVs: **(a)** Co-immunoprecipitation assay showing interaction of different domains of lactoferrin N protein with GAPDH. Different domains of lactoferrin N are shown in schematic at the top of the figure. Domains were fused to GFP and expressed in HEK293T cells. cMyc beads were used to capture GAPDH-cMyc fusion protein. The lactoferrin N2 domain showed interaction with GAPDH. Representative image (n>3). **(b)** Exogenous binding of GAPDH to EVs isolated from different cells. Higher molecular weight of exogenous GAPDH than endogenous GAPDH is due to presence of histidine tag (His6) and flag tag. **(c)** GAPDH kinetic assay of EVs before and after binding to exogenous GAPDH. Equal numbers of EVs were added to 96 well-plates in triplicates. The KDalert GAPDH assay kit was used to detect GAPDH activity. Results shown as mean ± s.d, n=3. **(d)** UV-absorbance spectrum of EVs after passing through gel-filtration column. Crude EVs present in the concentrated tissue culture medium were incubated with 1mg of GAPDH protein for 2h at 4°C and passed through the gel filtration column. Increase in the OD280 is due to binding of GAPDH to the EV surface and possibly aggregation of GAPDH-bound EVs. First peak (around 10ml elution) represents EVs and second peak (around 20 ml elution) represents tissue culture medium protein. Representative image (n>3).

**Extended Figure 3.**
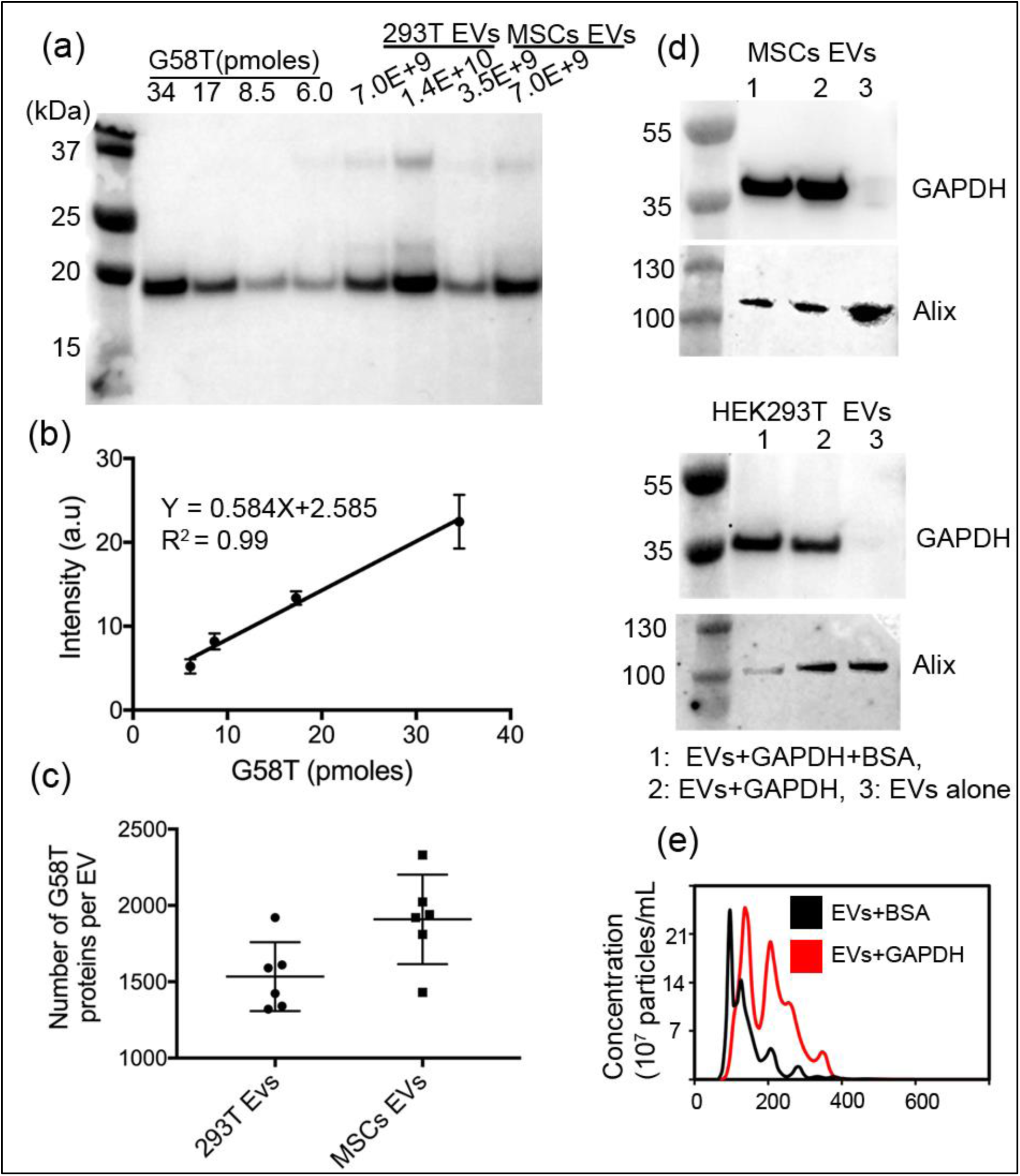
G58T quantification and *Drosophila* GAPDH binding to human EVs: **(a)**Western blot showing quantification of G58T protein on MSCs and HEK293T EVs (labelled as 293T EVs). 6.75 ×10^11^ EVs were incubated with 22 nmoles of G58T protein for 2h at 4°C. Excess of unbound protein was separated by gel-filtration chromatography. Specific amounts of purified G58T protein were run on SDS-PAGE along with EV samples to determine the number of G58T proteins on EVs. **(b)** Linear regression analysis of above immunoblots quantified using image studio software of *LI-COR*. **(c)**. Scatter plot showing number of G58T proteins bound to each MSC and 293T EV as determined by quantification of immunoblots. Data shown as mean ± s.d. n=3 (samples run in duplicates). **(d)** Western blots of MSC and 293T EVs after incubating with *Drosophila* GAPDH. 50ul of 2mg/mL BSA was added during the incubation. Representative blot (n=2). **(e)** Nanosight tracking analysis of EVs after incubation with *Drosophila* GAPDH protein. 20nmoles of GAPDH was incubated with 1.0 ×10^12^ EVs for 2h. A change in size of EVs after binding to GAPDH2 is consistent with clumping of EVs catalysed by *Drosophila* GAPDH. Representative graph (n=3).

**Extended Figure 4.**
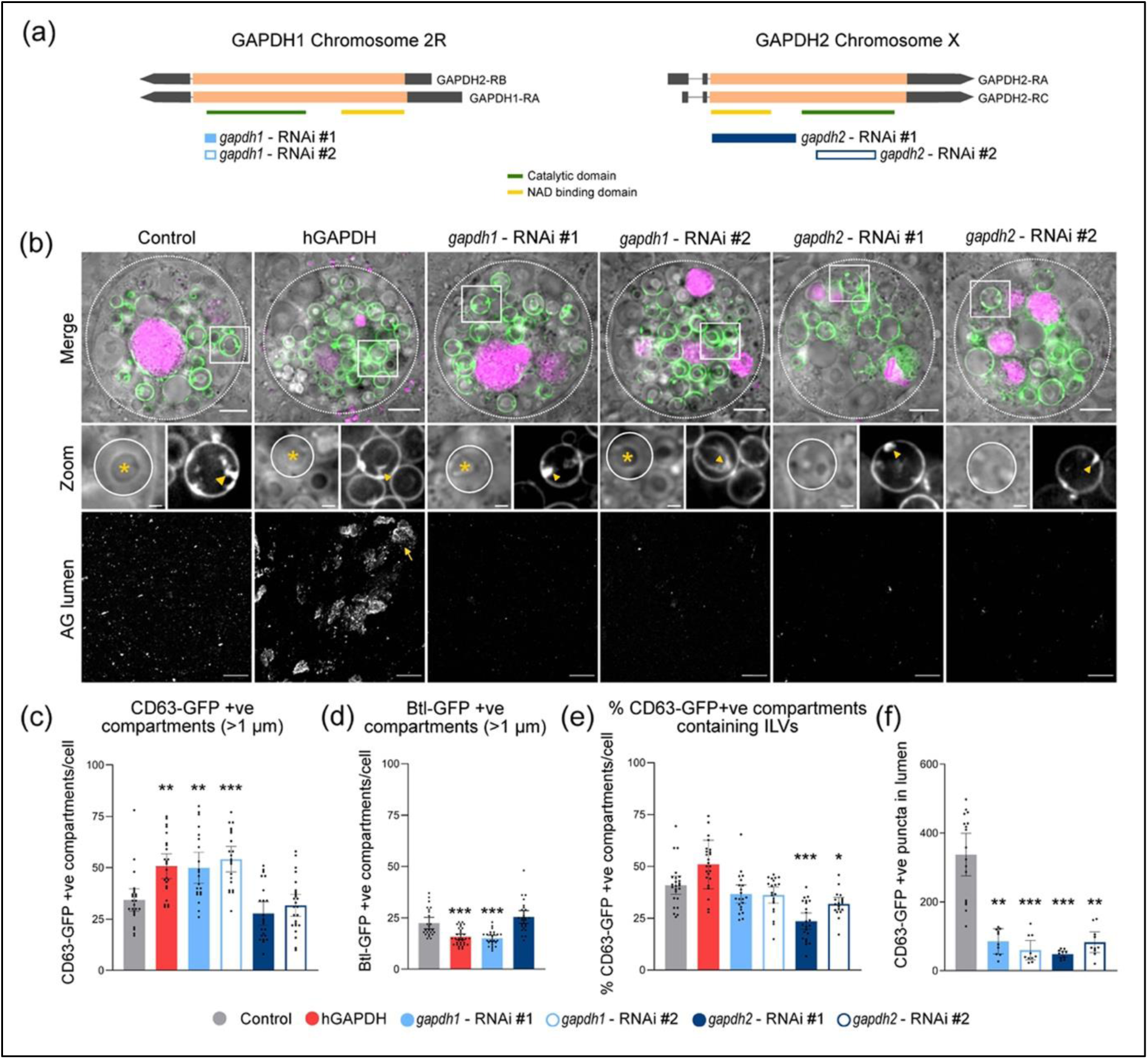
Manipulating GAPDH levels in *Drosophila* secondary cells affects the biogenesis of CD63-GFP-labelled exosomes: **(a)** Schematic shows isoforms of *Drosophila* GAPDH1 and GAPDH2 and the targeted regions of each RNAi line used. The two RNAi lines that target *GAPDH1* and do not have off-target effects bind to the same mRNA sequence. The two protein domains, the highly conserved catalytic domain and the NAD-binding domain, are also shown. **(b)** Basal wide-field fluorescence and differential interference contrast (‘Merge’) views of living secondary cells (SCs) expressing a GFP-tagged form of CD63 (CD63-GFP; green) with no other transgene (control); or also expressing the open reading frame of the human GAPDH protein (hGAPDH), independent RNAi constructs targeting *Drosophila* GAPDH1 (*gapdh1* – RNAi #1 and #2), or independent RNAi constructs targeting *Drosophila* GAPDH2 (*gapdh2* – RNAi #1 and #2). SC outline approximated by dashed white circles, and acidic compartments are marked by LysoTracker Red (magenta). CD63-GFP-positive intraluminal vesicles (ILVs; green in ‘Merge’; grey in ‘Zoom’) are apparent inside compartments, surrounding dense-core-granules (DCGs; asterisk in ‘Zoom’) and connecting DCGs to the limiting membrane of the compartment (yellow arrowheads, except in *GAPDH2* knockdown, where ILVs only surround peripheral small DCGs). DCG compartment outline is approximated by white circles. Confocal transverse images of fixed accessory gland (AG) lumens from the same genotypes, containing CD63-GFP fluorescent puncta. **(c)** Bar chart shows average number of large (> 1 µm) CD63-GFP-positive compartments per cell, which is increased following hGAPDH and *GAPDH1*-RNAi expression. CD63-GFP expression increases the number of non-acidic compartments when expressed by itself in SCs (Redhai et al., 2016 - PMID: 27727275). **(d)** Bar chart, relating to Figure 2, shows average number of large (> 1 µm) Btl-GFP-positive compartments per cell, which is slightly reduced following hGAPDH overexpression or *GAPDH1* knockdown. **(e)** Bar chart shows the percentage of CD63-GFP-positive compartments per cell containing ILVs. This number is selectively reduced by *GAPDH2* knockdown, which also reduces the normal clustering of ILVs in these compartments. **(f)** Bar chart shows CD63-GFP fluorescent puncta in the lumen of AGs for the different genotypes. Clustering of exosomes in the presence of hGAPDH prevented accurate quantification. All data are from six-day-old male flies shifted to 29°C at eclosion to induce expression of transgenes. Genotypes are: *w; P[w^+^, UAS-CD63-GFP] P[w^+^, tub-GAL80^ts^]/+*; *dsx-GAL4/+* with no other transgene (control), UAS-hGAPDH, UAS-*gapdh1*-RNAi #1 and #2, UAS-*gapdh2*-RNAi #1 and #2. Scale bars in (b) (5 µm), in ‘Zoom’ (1 µm), and in ‘AG lumen’ (20 µm). Data were analysed by the Kruskal-Wallis test followed by Dunn’s multiple comparison test. *** P <0.001, ** P < 0.01 and * P < 0.05 relative to control, n=23-27 cells, (f) n=10 glands.

**Extended Figure 5.**
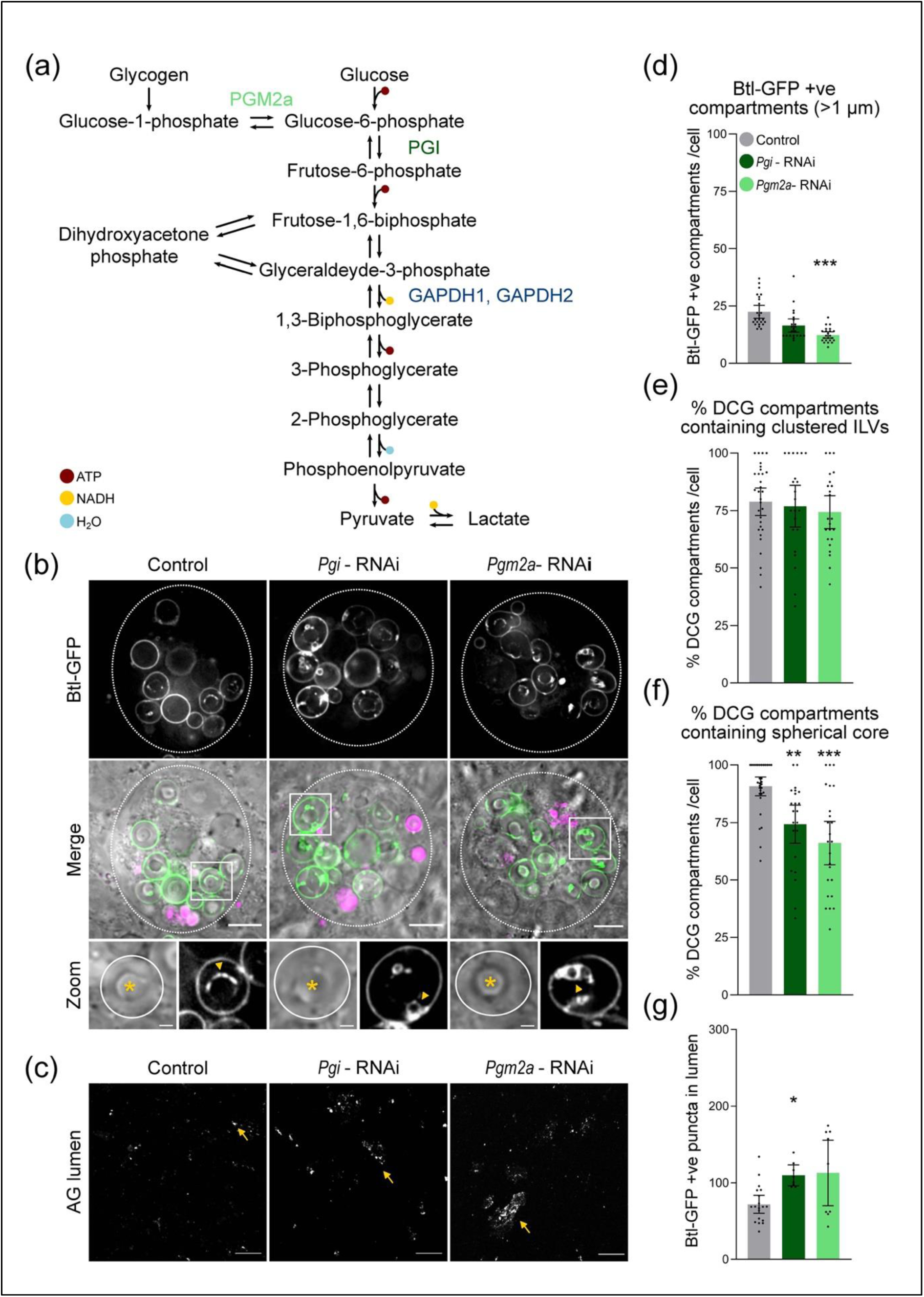
The effects of knocking down other glycolytic enzymes on exosome biogenesis and clustering in *Drosophila* secondary cells: **(a)** Schematic shows a simplified glycolytic pathway, in which the four glycolytic enzymes that were analysed in this study, GAPDH1 and 2, Phosphoglucomutase 2a (Pgm2a), and Phospho-glucose isomerase (Pgi) are highlighted. **(b)** Basal wide-field fluorescence and differential interference contrast (‘Merge’) views of living secondary cells (SCs) expressing a GFP-tagged form of Breathless (Btl-GFP; green) with no other transgene (control); or also expressing an RNAi construct targeting *Drosophila* Phosphoglucose isomerase (*Pgi*-RNAi), or *Drosophila* Phosphoglucomutase 2a (*Pgm2a*-RNAi). SC outline approximated by dashed white circles, and acidic compartments are marked by LysoTracker Red (magenta). Btl-GFP-positive intraluminal vesicles (ILVs; green in ‘Merge’; grey in ‘Zoom’) are apparent inside compartments (yellow arrowheads), surrounding dense-core-granules (DCGs; asterisk in ‘Zoom’). DCG compartment outline is approximated by white circles. **(c)** Confocal transverse images of fixed accessory gland (AG) lumens from the same genotypes, revealing Btl-GFP fluorescent puncta, which are more clustered in all knockdowns, perhaps because levels of GAPDH2 that are complexed with these enzymes are reduced. **(d)** Bar chart shows average number of large (> 1 µm) Btl-GFP-positive compartments per cell, which is slightly reduced in both knockdowns. **(e)** Bar chart shows percentage of DCG compartments per cell containing clustered Btl-GFP-positive ILVs that are not directly in contact with DCGs. This number is unaffected in both knockdowns in sharp contrast to *GAPDH2* knockdown (Fig 2). **(f)** Bar chart shows the percentage of DCG compartments per cell containing a single spherical core; a small number of abnormally shaped cores is observed in both knockdowns. **(g)** Bar chart shows number of Btl-GFP fluorescent puncta in the lumen of AGs from the same genotypes, which is unaffected or slightly increased in the knockdowns. All data are from six-day-old male flies shifted to 29°C at eclosion to induce expression of transgenes. Genotypes are: *w; P[w^+^, tub-GAL80^ts^]/+*; *dsx-GAL4, P[w^+^, UAS-btl-GFP]/+* with no other transgene (control), UAS-*Pgm2a*-RNAi, UAS-*Pgi*-RNAi or UAS-*Pfk*-RNAi. Scale bars in **(b)** (5 µm), in ‘Zoom’ (1 µm), and in **(c)** (20 µm). *** p <0.001, ** p < 0.01 and * p < 0.05 relative to control, n=22-30 cells, (g) n=8-10 glands.

**Extended Figure 6.**
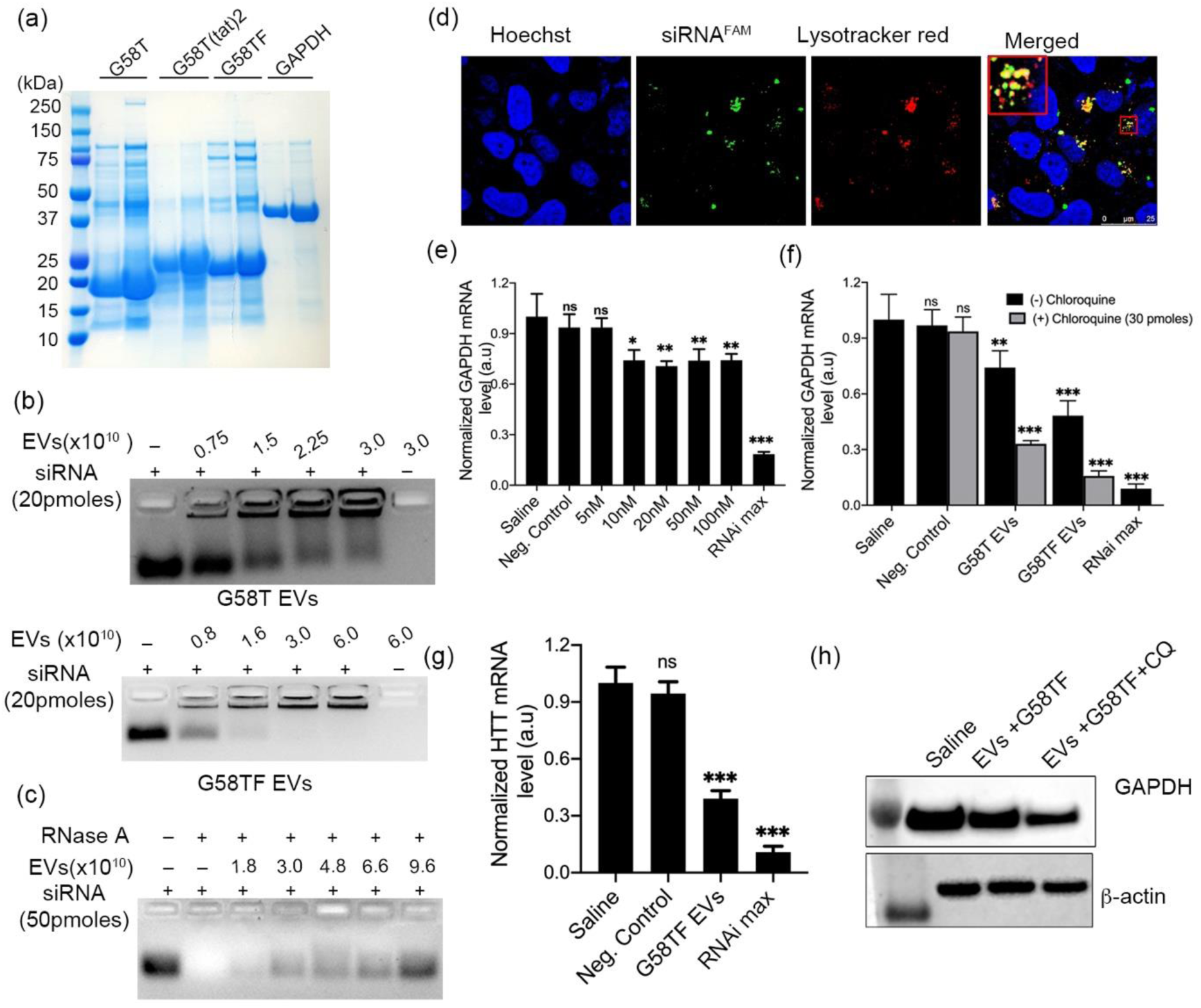
G58T-mediated loading of siRNA on EVs: SDS-PAGE of the G58T-based proteins purified by Ni-NTA chromatography. Gel is stained with Coomassie brilliant blue dye. **(b)** Gel shift assay of G58T EVs (upper) and G58TF EVs (lower), reflecting binding of siRNA to EVs. 20pmoles of siRNA were incubated with the given number of EVs. Bands trapped near the wells represent bound siRNA. siRNA alone was used as negative control. Based on the gel shift assay, around 550 siRNAs bind to each G58T EV and 714 siRNAs to each G58TF EV. Higher binding of siRNA to G58TF is due to the arginine-rich FHV peptide. **(c)** RNase A protection assay of G58TF EVs bound to 50 pmoles of siRNA. Bands in the agarose gel represent siRNA isolated from G58TF EVs after treatment with RNase A (0.2 mg/ml). G58TF EVs bound to siRNA were incubated with RNase A for 6 h at 37°C. **(d)** Representative confocal microscopy images of N2a cells after 4h of treatment with G58TF EVs carrying FAM-labelled siRNA (green). Cells were treated with lysotracker red dye (red dots) to label late-endosomes. Inset in the merged figure represents magnified image of N2a cells showing colocalization of siRNA with lysotracker dye (yellow spots). Images were captured using 60X objective lens of Olympus confocal microscope FV1000. **(e)** GAPDH gene silencing by G58T EVs at mRNA level in N2a cells. Different amounts of GAPDH siRNA bound to G58T EVs were added to cells. Level of mRNA was quantified after 48h of treatment by probe-based real time PCR. Data were analysed using linear regression analysis and ΔΔCt method. Level of mRNA in control was assigned as 1 and used to determine the relative level of GAPDH mRNA in the treated cells. Results are shown as mean ± s.d, n= 3, ***p < 0.0001 when compared to control (one-way ANOVA) **(f)** Graph showing effect of chloroquine on silencing of GAPDH by G58T and G58TF EVs in N2a cells. 30µM of Chloroquine was added during treatment of N2a cells with the EVs. Graph shows the level of GAPDH mRNA determined by probe-based real time PCR after 48h of treatment. Results are shown as mean ± s.d, n=3, **p =0.0047, ***P = 0.0001 when compared to control (one-way ANOVA). **(g)** Bar graph represents silencing of HTT gene in N2a cells after 48h of treatment with 50nM HTT siRNA bound to G58TF EVs. **(h)** Western blot showing GAPDH silencing in HeLa cells by G58TF EVs in presence and absence of chloroquine (30µM). 40pmoles of siRNA bound to EVs were added to cells and incubated for 72 h. Representative blot (n =2)

**Extended Figure 7.**
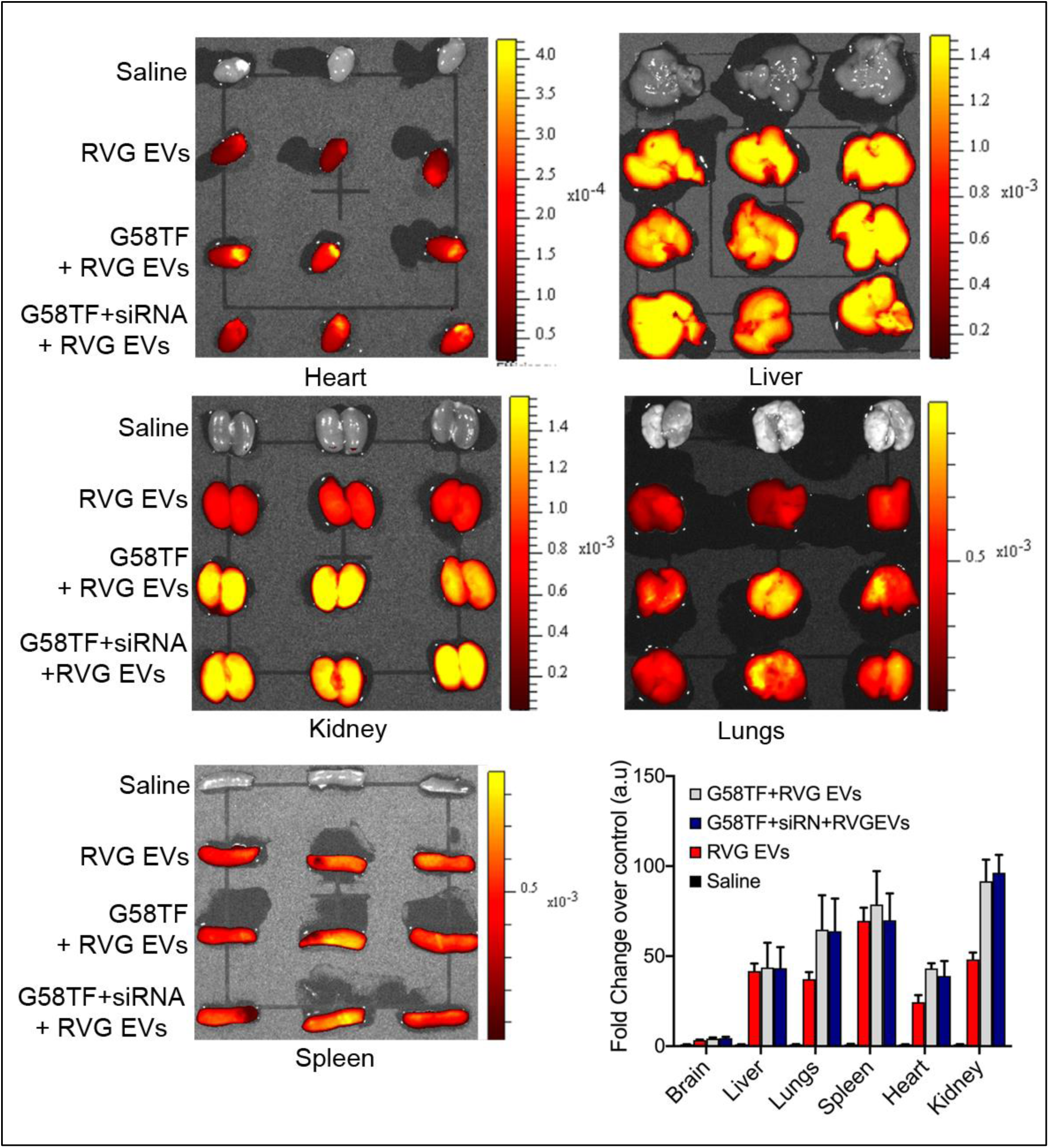
Biodistribution of systemically administered modified EVs. *In vivo* animal imaging of C57BL/6 animals after 4 h of administration of G58TF EVs labelled with cy-5.5 dye. Images represent fluorescence of cy-5.5 dye in various organs of the animals. Modification of EV surface with G58TF and siRNA did not alter biodistribution of RVG EVs. Three animals were used in each group. Graph shows quantification of the signals in different organs of the animals. Results are shown as mean ± s.d

## Supporting data

### Expression of a lactoferrin-LAMP-2B fusion protein on the surface of extracellular vesicles (EVs) results in cleavage but not release of lactoferrin from the EV surface

To facilitate targeted delivery of EVs into brain, we selected lactoferrin protein for expression on the EV surface. Lactoferrin is an iron-binding protein that is reported to cross the blood-brain barrier *via* receptor-mediated transcytosis^1,2^. The N-terminal domain of lactoferrin (designated as lactoferrin N) interacts with the lactoferrin receptor for internalization^3^. We fused lactoferrin N with lysosome-associated membrane protein of dendritic cells (DC-LAMP or LAMP3) to express it on the surface of EVs. During the expression, we noticed cleavage of lactoferrin N from DC-LAMP. Moreover, immunoblotting of HEK293T EVs revealed association of cleaved lactoferrin N with purified EVs **(Supporting Fig. 1)**. Attachment of green fluorescent protein (GFP) to DC-LAMP did not result in cleavage of GFP from DC-LAMP, suggesting that lactoferrin cleavage could be due to the presence of a specific protease cleavage site in the lactoferrin-DC-LAMP construct (**Supporting Fig. 2a, vector 1)**. Furthermore, attachment of lactoferrin N to other isoforms of LAMP proteins such as LAMP-2B (DC-LAMP contains only one luminal domain and LAMP-2B contains two luminal domains) did not change the nature of lactoferrin N cleavage, suggesting the specific cleavage site is in lactoferrin N region of the construct **(Supporting Fig. 2a, vector 2 &3)**. To determine the exact location of the cleavage site, we inserted cMyc and flag tags on either side of the linker sequence of lactoferrin-DC-LAMP (construct 4). Expression of construct 4 in cells confirmed presence of a specific proteolytic cleavage site in the lactoferrin region **(Supporting 2a &b)**. Bioinformatic analysis of the lactoferrin sequence, using Protease Specificity Prediction Server (PROSPER), revealed the presence of a specific group II cleavage site (L/IXX/Xhyd) of matrix metalloproteinase-2 (MMP-2) near the C-terminus of lactoferrin N, consistent with the molecular weight of cleaved lactoferrin N protein. A group IV cleavage site (XhydSX/L, S and L are invariant) of MMP-2 was also detected by PROSPER near the N-terminus of the lactoferrin **(Supporting Fig. 2c)**. However, lack of an important isoleucine renders it less prone to cleavage by MMP-2^4^. To assess whether MMP-2 is responsible for cleavage of the lactoferrin N, we mutated the group II cleavage site and fused a cMyc tag to the N-terminus of lactoferrin N. The protein expressed from this construct in HEK293T was not cleaved **(Supporting Fig. 2d)**. The full-length lactoferrin-LAMP-2B protein was detected with cMyc and lactoferrin antibodies indicating that MMP-2 also does not cleave the group IV site. Immunoblotting of HEK293T EVs revealed the association of MMP-2 with EVs **(Supplementary Fig. 2e).** Taken together, the data suggest cleavage of the protein by MMP-2 enzyme. Although it seems likely that cleavage is occurring on the surface of EVs during EV biogenesis, it remains possible that lactoferrin is cleaved in the cell cytoplasm, and later trafficked to the EV surface by GAPDH protein, which tethers cleaved lactoferrin to the surface of EVs, as explained in other parts of the manuscript.

**Supporting data Figure 1.**
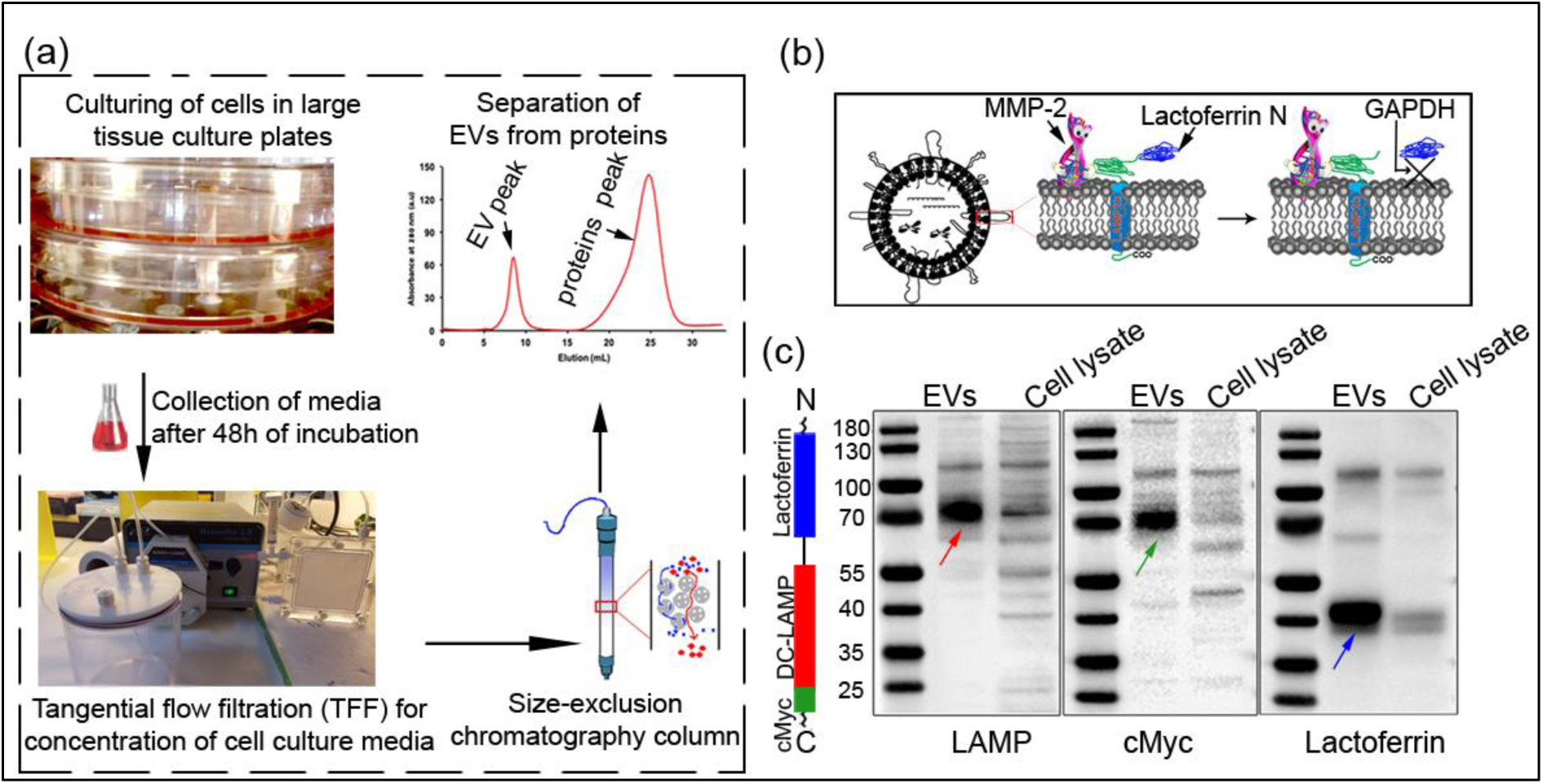
Association of proteolytically cleaved lactoferrin N with extracellular vesicles: **(a)** Schematic representation of EV-purification from cell culture medium. EVs eluted from gel-filtration column were used for further analysis. **(b)** A cartoon diagram showing cleavage of lactoferrin from DC-LAMP, and attachment of the cleaved lactoferrin on the EV surface via GAPDH protein. **(c)** Western blotting of purified HEK293T EVs and cell lysate showing expression of lactoferrin N-DC-LAMP. Three antibodies detecting LAMP, cMyc and lactoferrin were used to assess the expression. Detection of the lactoferrin N band at lower position (blue arrow) than the LAMP (Red arrow) and cMyc band (green arrow) reveals cleavage of lactoferrin N from DC-LAMP. Arrangement of the different domains of the protein is shown on the left side of the blot. Presence of cleaved lactoferrin N in the EV sample indicates association of the protein with EVs. Representative blot (n>3).

**Supporting data Figure 2.**
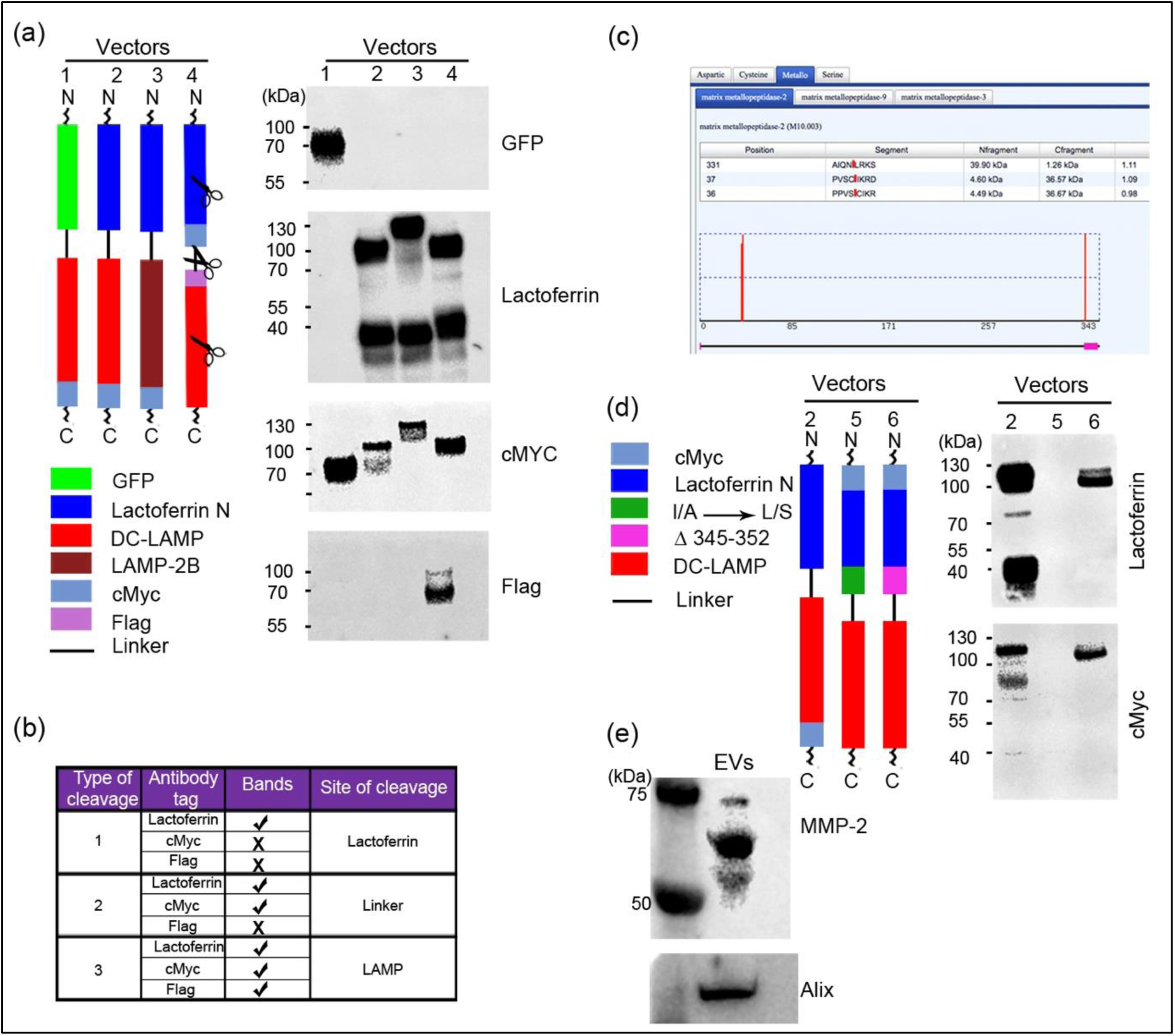
Matrix metalloproteinase-2 is involved in proteolytic cleavage of lactoferrin N: **(a & b)** Design and expression of various lactoferrin-LAMP constructs in HEK293T cells. Left side of the figure is a schematic diagram of the vector constructs. Expression of these constructs in HEK293T is shown by western blotting of HEK293T cell lysates using respective antibodies. Vector 1 expresses GFP-DC-LAMP. A single band detected at the same position by GFP and cMyc antibodies confirms intact GFP-DC-LAMP chimeric protein in the cells. Vector 2 and 3 express lactoferrin N attached to either DC-LAMP or LAMP-2B protein respectively. Expression of both the vectors resulted in cleavage of lactoferrin N domain (two bands in lactoferrin blot). In vector 4, cMyc and Flag tags was inserted on either side of the linker sequence to identify the cleavage site. Table **(b)** shows three different types of cleavage patterns of vector 4 depending upon the location of cleavage site. Expression of vector 4 in HEK293T cells showed cMyc and flag tag bands (lane marked with 4 in (a)) at same position and no bands for either tags were detected near cleaved lactoferrin N position, confirming presence of cleavage site in the lactoferrin N domain. **(c)** *In silico* prediction of protease cleavage sites in the lactoferrin N domain. Protease Specificity Prediction Server (PROSPER) was used to determine the possible protease cleavage sites (https://prosper.erc.monash.edu.au). The data showed a highly specific group II MMP-2 cleavage site at amino acid 331 and a group IV cleavage site at amino acid 37. **(d)** Design and expression of mutated lactoferrin-LAMP construct, lacking MMP-2 cleavage II site. Vector 2 was used as a positive control for MMP-2 cleavage. In vector 5, specific residues were mutated to inactivate the MMP-2 cleavage site, but the protein was not detected by the lactoferrin antibody for reasons that are unclear. In vector 6, the group II cleavage site was completely deleted, which results in the absence of the lower lactoferrin N band, consistent with MMP-2 mediated cleavage of lactoferrin N. Detection of a single band by cMyc and lactoferrin antibodies in vector 6 reflects the inability of MMP-2 to cleave at the group IV cleavage site present at the N-terminus of lactoferrin. **(e)** Western blotting of HEK293T EVs showing expression of MMP-2 enzyme. All experiments were carried out in triplicate in 6-well tissue culture plates. Vectors were transiently expressed in the cells using standard transfection protocols.

**Supporting data Figure 3:**
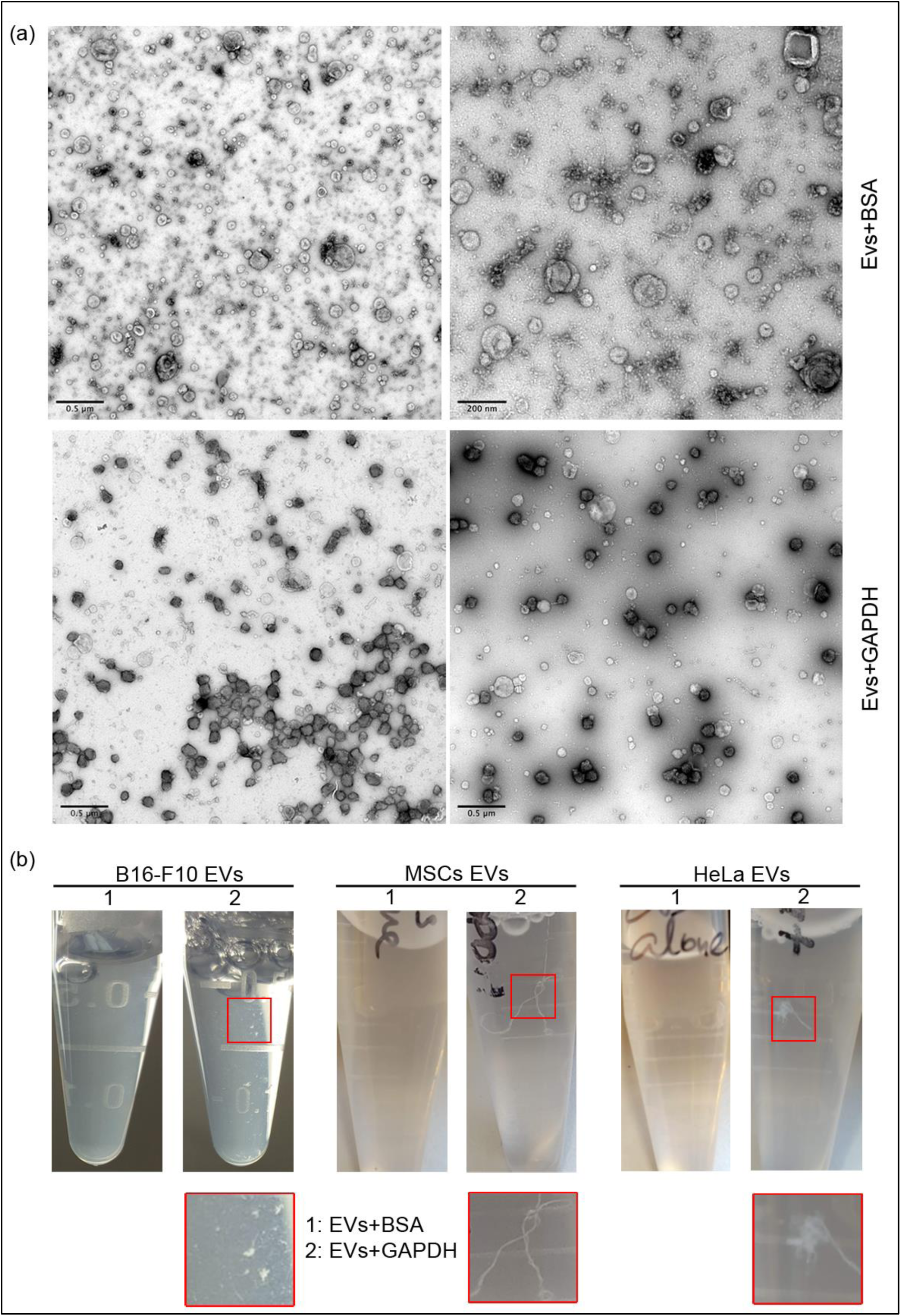
**(a)** Electron microscopy (EM) images of HEK293T extracellular vesicles after incubation with either BSA or GAPDH protein. Excess of unbound proteins was separated prior to EM imaging. The images show aggregation of EVs mediated by GAPDH protein. **(b)** Photographic images of the EVs after 2h of addition of either BSA or GAPDH protein. Addition of GAPDH lead to formation of conspicuous EV aggregates. Figure (a) and (b) represent raw images of Figure (1d) and (1e) respectively.

**Supporting data Figure 4:**
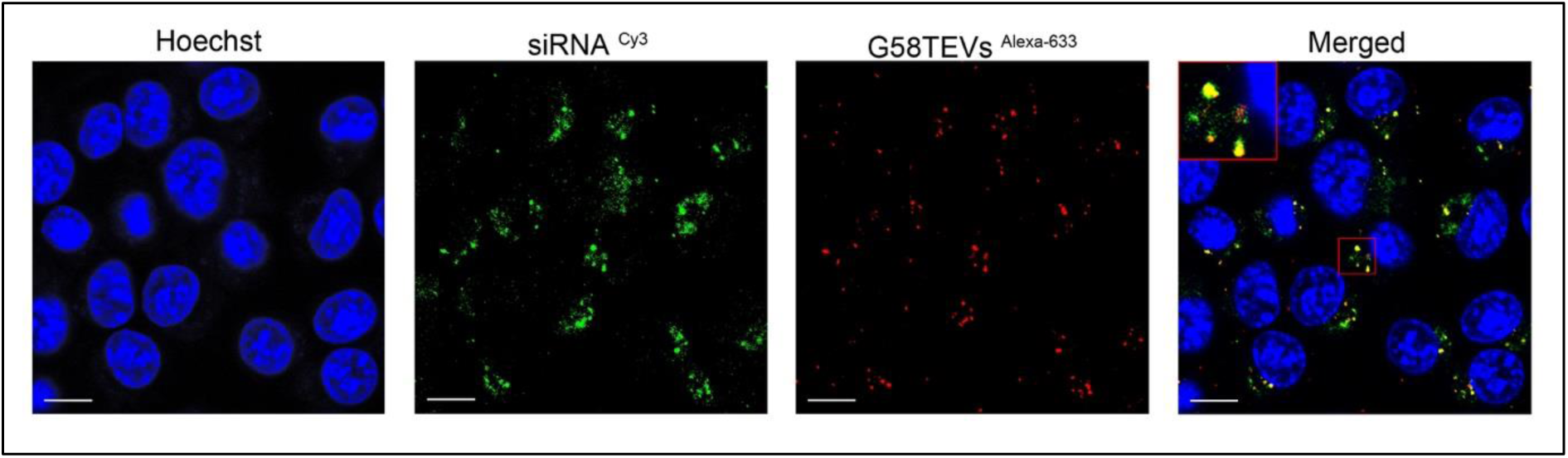
Confocal microscopy images of N2a cells showing uptake of G58T EVs carrying siRNA. Prior to loading of G58T protein, surface proteins of EVs were conjugated with Alexa Fluor-633 (red dots). siRNA loaded into EVs was conjugated with Cy-3 fluorescent dye (pseudo-green colour). Images were captured using 60X objective lens of Olympus confocal microscope FV1000. Representative images (n>3).

## References

1 Raposo, G. & Stahl, P. D. Extracellular vesicles: a new communication paradigm? Nat Rev Mol Cell Biol 20, 509–510, doi:10.1038/s41580-019-0158-7 (2019).

2 Tkach, M. & Thery, C. Communication by Extracellular Vesicles: Where We Are and Where We Need to Go. Cell 164, 1226–1232, doi:10.1016/j.cell.2016.01.043 (2016).

3 Willms, E., Cabanas, C., Mager, I., Wood, M. J. A. & Vader, P. Extracellular Vesicle Heterogeneity: Subpopulations, Isolation Techniques, and Diverse Functions in Cancer Progression. Front Immunol 9, 738, doi:10.3389/fimmu.2018.00738 (2018).

4 S, E. L. A., Mager, I., Breakefield, X. O. & Wood, M. J. Extracellular vesicles: biology and emerging therapeutic opportunities. Nat Rev Drug Discov 12, 347–357, doi:10.1038/nrd3978 (2013).

5 Johnsen, K. B. et al. A comprehensive overview of exosomes as drug delivery vehicles - endogenous nanocarriers for targeted cancer therapy. Biochim Biophys Acta 1846, 75–87, doi:10.1016/j.bbcan.2014.04.005 (2014).

6 Sterzenbach, U. et al. Engineered Exosomes as Vehicles for Biologically Active Proteins. Mol Ther 25, 1269–1278, doi:10.1016/j.ymthe.2017.03.030 (2017).

7 Tang, T. T., Lv, L. L., Lan, H. Y. & Liu, B. C. Extracellular Vesicles: Opportunities and Challenges for the Treatment of Renal Diseases. Front Physiol 10, 226, doi:10.3389/fphys.2019.00226 (2019).

8 Colombo, M., Raposo, G. & Thery, C. Biogenesis, secretion, and intercellular interactions of exosomes and other extracellular vesicles. Annu Rev Cell Dev Biol 30, 255–289, doi:10.1146/annurev-cellbio-101512-122326 (2014).

9 Kooijmans, S. A. A. et al. Electroporation-induced siRNA precipitation obscures the efficiency of siRNA loading into extracellular vesicles. J Control Release 172, 229–238, doi:10.1016/j.jconrel.2013.08.014 (2013).

10 Smyth, T. et al. Surface functionalization of exosomes using click chemistry. Bioconjug Chem 25, 1777–1784, doi:10.1021/bc500291r (2014).

11 Sutaria, D. S., Badawi, M., Phelps, M. A. & Schmittgen, T. D. Achieving the Promise of Therapeutic Extracellular Vesicles: The Devil is in Details of Therapeutic Loading. Pharm Res 34, 1053–1066, doi:10.1007/s11095-017-2123-5 (2017).

12 Murphy, D. E. et al. Extracellular vesicle-based therapeutics: natural versus engineered targeting and trafficking. Exp Mol Med 51, 32, doi:10.1038/s12276-019-0223-5 (2019).

13 Malhotra, H. et al. Exosomes: Tunable Nano Vehicles for Macromolecular Delivery of Transferrin and Lactoferrin to Specific Intracellular Compartment. J Biomed Nanotechnol 12, 1101–1114, doi:10.1166/jbn.2016.2229 (2016).

14 Glaser, P. E. & Gross, R. W. Rapid plasmenylethanolamine-selective fusion of membrane bilayers catalyzed by an isoform of glyceraldehyde-3-phosphate dehydrogenase: discrimination between glycolytic and fusogenic roles of individual isoforms. Biochemistry 34, 12193–12203, doi:10.1021/bi00038a013 (1995).

15 Han, X., Ramanadham, S., Turk, J. & Gross, R. W. Reconstitution of membrane fusion between pancreatic islet secretory granules and plasma membranes: catalysis by a protein constituent recognized by monoclonal antibodies directed against glyceraldehyde-3-phosphate dehydrogenase. Biochim Biophys Acta 1414, 95–107, doi:10.1016/s0005-2736(98)00154-0 (1998).

16 Nakagawa, T. et al. Participation of a fusogenic protein, glyceraldehyde-3-phosphate dehydrogenase, in nuclear membrane assembly. J Biol Chem 278, 20395–20404, doi:10.1074/jbc.M210824200 (2003).

17 Kaneda, M., Takeuchi, K., Inoue, K. & Umeda, M. Localization of the phosphatidylserine-binding site of glyceraldehyde-3-phosphate dehydrogenase responsible for membrane fusion. J Biochem 122, 1233–1240, doi:10.1093/oxfordjournals.jbchem.a021886 (1997).

18 Corrigan, L. et al. BMP-regulated exosomes from Drosophila male reproductive glands reprogram female behavior. J Cell Biol 206, 671–688, doi:10.1083/jcb.201401072 (2014).

19 Fan, S.-J. et al. Glutamine deprivation regulates the origin and function of cancer cell exosomes. BioRxiv, https://doi.org/10.1101/859447.

20 Dar, G. H., Gopal, V. & Rao, M. Conformation-dependent binding and tumor-targeted delivery of siRNA by a designed TRBP2: Affibody fusion protein. Nanomedicine 11, 1455–1466, doi:10.1016/j.nano.2015.01.017 (2015).

21 Wadia, J. S., Stan, R. V. & Dowdy, S. F. Transducible TAT-HA fusogenic peptide enhances escape of TAT-fusion proteins after lipid raft macropinocytosis. Nat Med 10, 310–315, doi:10.1038/nm996 (2004).

22 Worch, R. Structural biology of the influenza virus fusion peptide. Acta Biochim Pol 61, 421–426 (2014).

23 Nakase, I. et al. Cell-surface accumulation of flock house virus-derived peptide leads to efficient internalization via macropinocytosis. Mol Ther 17, 1868–1876, doi:10.1038/mt.2009.192 (2009).

24 Wolfert, M. A. & Seymour, L. W. Chloroquine and amphipathic peptide helices show synergistic transfection in vitro. Gene Ther 5, 409–414, doi:10.1038/sj.gt.3300606 (1998).

25 Godinho, B. et al. Transvascular Delivery of Hydrophobically Modified siRNAs: Gene Silencing in the Rat Brain upon Disruption of the Blood-Brain Barrier. Mol Ther 26, 2580–2591, doi:10.1016/j.ymthe.2018.08.005 (2018).

26 Alvarez-Erviti, L. et al. Delivery of siRNA to the mouse brain by systemic injection of targeted exosomes. Nat Biotechnol 29, 341–345, doi:10.1038/nbt.1807 (2011).

27 Wiklander, O. P. et al. Extracellular vesicle in vivo biodistribution is determined by cell source, route of administration and targeting. J Extracell Vesicles 4, 26316, doi:10.3402/jev.v4.26316 (2015).

28 Glorioso, J. C., Cohen, J. B., Carlisle, D. L., Munoz-Sanjuan, I. & Friedlander, R. M. Moving toward a gene therapy for Huntington’s disease. Gene Ther 22, 931–933, doi:10.1038/gt.2015.102 (2015).

29 Menalled, L. B., Sison, J. D., Dragatsis, I., Zeitlin, S. & Chesselet, M. F. Time course of early motor and neuropathological anomalies in a knock-in mouse model of Huntington’s disease with 140 CAG repeats. J Comp Neurol 465, 11–26, doi:10.1002/cne.10776 (2003).

30 Kurosawa, M. et al. Depletion of p62 reduces nuclear inclusions and paradoxically ameliorates disease phenotypes in Huntington’s model mice. Hum Mol Genet 24, 1092–1105, doi:10.1093/hmg/ddu522 (2015).

31 Nordin, J.Z. et al. Ultrafiltration with size-exclusion liquid chromatography for high yield isolation of extracellular vesicles preserving intact biophysical and functional properties, Nanomedicine 11, 879–83 doi: 10.1016/j.nano.2015.01.003 (2015).

32 Théry, C., Amigorena, S., Raposo, G., & Clayton, A.. Isolation and characterization of exosomes from cell culture supernatants and biological fluids. Curr Protoc Cell Biol. Chapter 3:Unit 3.22. doi: 10.1002/0471143030.cb0322s30 (2006).

33. Nakase, I. et al. Cell-surface accumulation of flock house virus-derived peptide leads to efficient internalization via macropinocytosis. Mol. Ther. 17, 1868–76. doi: 10.1038/mt.2009.192 (2009).

34. Panáková, D., Sprong, H., Marois, E., Thiele C., and S. Eaton, S. Lipoprotein particles are required for Hedgehog and Wingless signalling. Nature 435, 58–65. doi: 10.1038/nature03504 (2005).

35. Rideout, E.J., Dornan, A.J., Neville, M.C., Eadie, S., & Goodwin, S.F. Control of sexual differentiation and behavior by the *doublesex* gene in *Drosophila melanogaster*. Nat. Neurosci. 13, 458–466. doi: 10.1038/nn.2515 (2010).

36. Sato, M., & Kornberg, T.B. FGF is an essential mitogen and chemoattractant for the air sacs of the Drosophila tracheal system. Dev. Cell 3, 195–207. doi: 10.1016/s1534-5807(02)00202-2 (2002).

37. Perkins, L. A., et al. The Transgenic RNAi Project at Harvard Medical School: Resources and Validation. Genetics 201, 843–52. doi: 10.1534/genetics.115.180208 (2015).

38. Dietzl, G., et al. A genome-wide transgenic RNAi library for conditional gene inactivation in *Drosophila*. Nature 448, 151–156. doi: 10.1038/nature05954 (2007).

39. Leiblich, A., et al. Bone morphogenetic protein- and mating-dependent secretory cell growth and migration in the *Drosophila* accessory gland. Proc Natl Acad Sci USA. 109, 19292–7. doi: 10.1073/pnas.1214517109 (2012).

40. Parton, R.M., Vallés, A.M., Dobbie, I.M., & Davis, I. Live Cell Imaging in *Drosophila melanogaster*. Cold Spring Harb. Protoc. doi:10.1101/pdb.top75 (2010).

41. Schindelin, J., Rueden, C.T., Hiner, M.C., & Eliceiri, K.W. The ImageJ ecosystem: an open platform for biomedical image analysis. Mol. Reprod. Dev. 82, 518–529. doi: 10.1002/mrd.22489 (2015)

## References

1 Huang, R. Q. et al. Characterization of lactoferrin receptor in brain endothelial capillary cells and mouse brain. J Biomed Sci 14, 121–128, doi:10.1007/s11373-006-9121-7 (2007).

2 Fillebeen, C. et al. Receptor-mediated transcytosis of lactoferrin through the blood-brain barrier. J Biol Chem 274, 7011–7017, doi:10.1074/jbc.274.11.7011 (1999).

3 Suzuki, Y. A., Wong, H., Ashida, K. Y., Schryvers, A. B. & Lonnerdal, B. The N1 domain of human lactoferrin is required for internalization by caco-2 cells and targeting to the nucleus. Biochemistry 47, 10915–10920, doi:10.1021/bi8012164 (2008).

4 Chen, E. I. et al. A unique substrate recognition profile for matrix metalloproteinase-2. J Biol Chem 277, 4485–4491, doi:10.1074/jbc.M109469200 (2002).

